# Rhizosphere metagenomics of mine tailings colonizing plants: assembling and selecting synthetic bacterial communities to enhance *in situ* bioremediation

**DOI:** 10.1101/664805

**Authors:** Miguel Romero, Diana Gallego, Jazmin Blaz, Arely Lechuga, José F. Martínez, Hugo R. Barajas, Corina Hayano-Kanashiro, Mariana Peimbert, Rocío Cruz-Ortega, Francisco E. Molina-Freaner, Luis D. Alcaraz

## Abstract

Mine tailings phytostabilization has been proposed as a bioremediation strategy to constrain the contaminants dispersion using plants to limit the effects of erosion. Rhizospheric bacteria impact plant health and facilitate plant establishment through their metabolic functions, which could be relevant in bioremediation strategies. We studied both culturable and metagenomic diversity or rhizospheric bacteria of mine tailings colonizing plants of an abandoned mine operation in Nacozari de García, Sonora, Mexico. Diversity was described through amplification of the 16S rRNA gene and whole metagenome shotgun sequencing of both environmental and cultured rhizosphere associated microbes. The culturable bacteria were assembled in a synthetic community (SC; 235 bacteria genera). Subsequently, we performed an experimental evolution setup with the SC, selecting for heavy metal resistance, microbial competition, and the ability for growing in plant-derived nutrient sources. The selection experiment show that bacteria diversity decreases from the environmental culture-free microbiomes to the mine tailings rhizospheres and the experimental evolution outcome: the synthetic community (FSC; 43 bacteria genera). The rhizosphere communities shifted from the dominance of *Actinobacteria* in their environment to *Proteobacteria* in the cultivated consortia and the synthetic communities. Both environmental and cultured metagenomes contained plant-growth promotion, heavy-metal homeostasis, and antibiotic resistance predicted genes. The FSC included predicted proteins related to plant-growth promotion such as siderophore production and plant hormone regulation proteins. We reconstructed a metagenome assembled genomic sequence named *Enterobacter* sp., Nacozari. The recovered *Enterobacter* sp. Nacozari, have predicted coding genes for direct and indirect plant growth promotion along with adhesion and oxidative stress-related proteins. The metabolic potential of the FSC presents promising features that might make it useful for plant-growth promotion in tailored phytostabilization strategies for the abandoned mine-tailings of Nacozari.

## Introduction

Unconfined mine tailings are harsh environments for plant colonization due to their conditions such as extreme pH, high concentrations of heavy metals, reduced water retention capacity, and autotrophic bacteria. In the absence of plants that reduce erosion, in arid and semi-arid climate regions these toxic wastes are dispersed through water and air currents that are dispersed to adjacent areas and are associated with human diseases (Meza-Figueroa *et al*.,2009; Nriagu 1988). Mine reclamation should be the ideal condition at the end of the mine operation, however, there is no regulation for old abandoned mines in multiple developing countries (Favas *et al*.,2018). Open mining reclamation strategies include phytostabilization which aims to reduce pollutants motility through plant coverage, ideally a plant community able to immobilize the metals in the roots, low shoot translocation, while recruiting heterotrophic bacteria and stabilizing the wasteland (Ali, Khan & Sajad, 2013).

A mine bioremediation *in situ* strategy known as phytostabilization, which prevent the exposure of mine waste into the environment, due to the use of local plants to prevent erosion through root consolidation of the mine tailings (Mendez & Maier 2008). Additionally, the plant roots exudates along the bacteria rhizosphere community precipitate heavy metals (Santos *et al*.,2017). A plant species is a candidate for phytostabilization of heavy metals if they are not translocated to shoots, keeping most of it precipitated or root-accumulated, thus preventing incorporation of heavy metals into trophic networks (*i*.*e*., cattle foraging) (Santos *et al*.,2017). In previous work, a spontaneously plant community colonization of an abandoned mine operation (*ca*. 1949) in north-western Mexico (Nacozari de García, Sonora, Mexico) was described (Santos *et al*.,2017). It was possible to identify plant species with bioremediation potential for the Nacozari mine tailings: *Acacia farnesiana, Brickellia coulteri, Baccharis sarothroides*, and *Gnaphalium leucocephalum* (Santos *et al*.,2017). Here we are following up that work and studying the microbial communities that could lead or restrict plant establishment in this location.

Previous work, has demonstrated that bacteria can be trained, as it has been done in the ongoing long term evolution experiment (LTEE), in which 12 *Escherichia coli* cell lines have been cultured in liquid glucose-limited medium and put through a bottleneck of 1% of the whole population in each serial pass into fresh medium each 24 hours. After the first 2,000 generations, the populations presented a higher fitness than the original cell lines growing in glucose-limited conditions (Lensky & Travisano, 1994), the experiment is now over 60,000 generations and it has been suggested that the fitness peak is not yet achieved (Good *et al*.,2017). Bacterial communities harbor different interactions such as cooperation, inhibition, and competition, while some members of the community might remain neutral and avoid all of the above resulting in community-intrinsic properties or properties of bacteria that are shown only in community level (Madsen *et al*.,2018). It has been shown that if a new member is introduced in the community, it will likely be sensitive to the antagonism effectors of the original members (Pérez-Gutiérrez *et al*., 2013). The build of synthetic communities is the scaling up of experimental evolution from single or two species interaction to integrating emergent member and molecular interactions of complex model systems, yet culturable and thus capable of testing experimentally the models, from genome interactions to predictable outputs (Cairns *et al*., 2018; Zomorrodi & Segrè 2016).

Plant-microbe interactions are critical for plant establishment, nutrient acquisition, and microbe interactions including neutral, parasitic, or mutualistic interactions affecting plants health (Bulgarelli *et al*.,2013). The plant roots-microbe interactions are essential for the establishment of the plant microbiome, some members of the bacteria community are capable of inducing plant-growth promotion (PGPB). PGPB comprehends multiple strategies, like nitrogen fixation, phosphate acquisition, plant-hormone production. For example, phosphate solubilization through the synthesis of organic acids and the production of phytases, mediated by the PqqBCDEFG proteins and AppA/A2, respectively (Liu *et al*.,1992; Rodriguez *et al*.,1999). In addition, rhizospheric bacteria can also modulate plant hormone concentrations by the synthesis of indole-3-acetic acid (IAA) with the enzyme phenylpyruvate decarboxylase, thus promoting root elongation (Spaepen *et al*.,2007) or by the consumption of 1-aminocyclopropane-1-carboxylate (ACC), an ethylene precursor, through the enzyme ACC deaminase, which maintain healthy ethylene concentrations in plant tissue (Glick *et al*.,1998). It has also been observed that ethylene pathways could help prevent salinity stress in plants, via bacterial production of ethylene precursors like 2-keto-4-methylthiobutyric acid (KMBA) (de Zélicourt, *et al*.,2018). Bacteria are also capable of immobilizing heavy metals through biosorption and dissimilatory reduction (Valls & De Lorenzo, 2002) while modulating their micronutrient concentration homeostasis through cation efflux pumps (Nies, 1999). Biofertilizers have been widely proposed as a sustainable and eco-friendly approach to promote plant growth (Bhardwaj et al.,2014). However, single species biofertilizers lack the community interactions described before, and it has been reported that simple communities (*≤*3 bacteria species) synergistic effects could lead to effective bioaugmentation and improvements in plant growth (Mansotra *et al*.,2015). Therefore, the design of biofertilizers inoculum should try to include entire communities. The design of synthetic bacterial communities is made with the rationale of harnessing the microbiome influence in the host phenotypes, and it is a newly developing field in the study of plant-microbe interactions (Herrera Paredes *et al*.,2018).

In this study, we describe both culturable and culture-free microbiomes from Nacozari mine-tailing rhizosphere microbe communities through 16S rRNA gene amplicons and whole shotgun metagenomic sequencing. We are introducing a mix of rhizospheric culturable strains into a synthetic community (SC), and then performing an experimental evolution setup looking for a culturable microbial community capable of surviving under mine-tailing conditions, also selected to depend solely on carbon sources derived from roots, and interact with the other bacteria community members (Figure 1).

**Figure 1.**
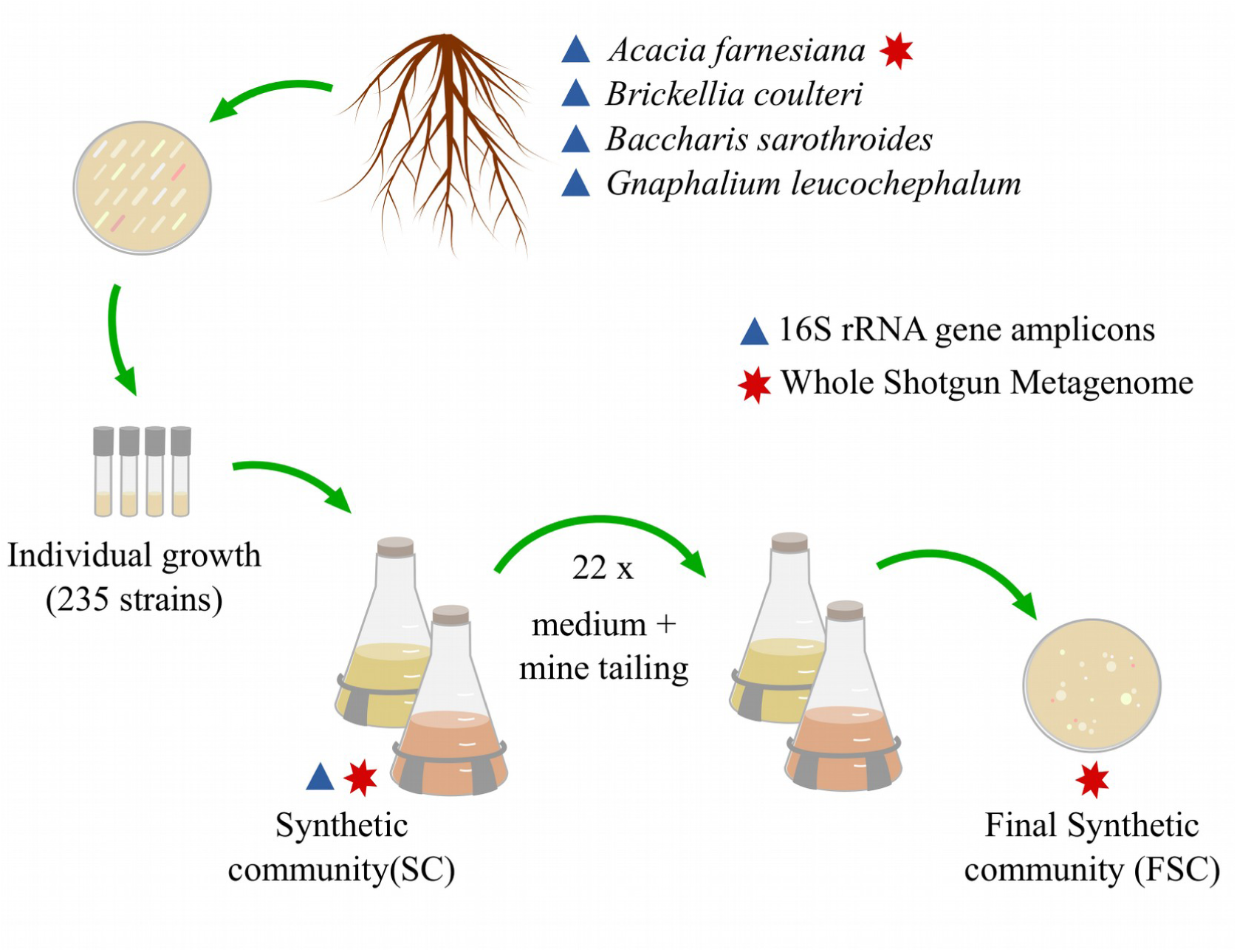
Overview of this work. *In situ* biodiversity of the Nacozari mine tailings was described both by 16S rRNA gene amplicons and shotgun sequencing. A total of 235 rhizosphere-derived bacterial strains were selected to assembly the synthetic community (SC). The SC was cultured in plant derived culture media or complex sugars, with an inhibiting mine tailings concentration for heavy-metal unadapted bacteria (see Material and Methods). Twenty-two serial passes of the SC were performed each 48 hours, the last serial pass is known as final synthetic community (FSC). The metabolic and taxonomic profiles were predicted for the SC and the FSC through the analysis of their shotgun metagenomes.

## Material and Methods

### Sampling

The town of Nacozari de García, Sonora is a mining district in northern Mexico (30°22′2.4″N, 109°41′38″O), it has a semi-arid climate with an annual mean temperature of 19.6°C and annual mean precipitation of 490 mm. The mine-tailing dam which lies adjacent to the residential zone (Supplementary Figure 1) contains quartz (SiO2), gypsum (CaSO4H2O), lepidocrocite (FeO[OH]) and copper sulfate (CuSO4, Romero et al., 2008). Mine tailing mean pH is 3.8 ± 0.3, and its average electrical conductivity is 340.1 ± 2 µS/cm (Meza-Figueroa et al., 2009). Root samples for amplicon and metagenomic DNA sequencing were collected on December, 1st, 2015, between 11:09 AM and 2:19 PM, for each plant species of *A. farnesiana* (Afp1), *B. coulteri* (Bc), *B. sarothroides* (Bs), and *G. leucocephalum* (Gl) growing on mine tailing, including an additional sample from an *A. farnesiana* plant growing in a secondary vegetation patch (Afp2; Supplementary Figure S1). Samples for metagenomic DNA extractions were stored in 50 ml sterile conical tubes and immediately stored in liquid nitrogen then stored in a freezer at −80 °C until DNA extraction. Sampling for microbiological cultures were collected on March 29th, 2016, from 7:26 AM to 8:20 PM. Duplicated root samples from plants species *A. farnesiana* (Afp1-CC), *B. coulteri* (Bc-CC), *B. sarothroides* (Bs-CC), and *G. leucocephalum* (Gl-CC) from the main tailing vegetation patch and an additional sample from an *A. farnesiana* (Afp2-CC) plant growing in a secondary vegetation patch (Supplementary Figure S1), were collected as described above, with the exception that the 50 ml sampling tubes containing 30 ml of sterile PBS solution and stored at 4°C until its processing.

### Microbiological procedures

The sampled tubes from plant rhizospheres were shaken and serially diluted; dilutions 10^−3^ and 10^−5^ were cultured in solid Luria Broth (LB) and Peptone Yeast (PY) media. The Petri dishes were then incubated (28°C). Single colonies were isolated, and colony forming units (cfu) were estimated from colony counts with averages of 10^6^ cfu. Isolated colonies were screened for the following morphological traits: form, border, elevation, surface, consistency, color, transmitted, and reflected light (Supplementary Figure S2). Results from the morphological variables were used as input to a multiple classification analysis (MCA) ordination to select representative colony diversity resulting in the selection of 235 isolates. Each isolate was grown in LB liquid media for 42. Absorbance was followed at 600 nm in a Genesys 20 spectrophotometer (Thermo Scientific). Each isolate was classified as slow, medium, and fast grower (Supplementary Figure S3).

### Experimental evolution setup

The 235 representative strains of bacteria morphological diversity were chosen as input for an experimental evolution mesocosm. We denominated this 235 strains subset as the synthetic community (SC). The goal was to assemble a synthetic community capable of tolerating mine tailing heavy metals, use plant-derived or complex carbon sources as main building block for their metabolism, and being capable to establish as a community and thus relationships of mutualism, and competition. Each one of the 235 isolates of the SC was grown until reaching an OD_600_ ∼ 0.4 and then 100 µL of culture transferred to fresh LB liquid media used all of them as the source for the experimental evolution setup.

We determined the concentration (w/v %) of the Nacozari mine tailings (Agate mortar ground, 2 rounds of autoclave sterilization) capable of inhibiting growth of two type strains *Escherichia coli* BW25113 and *Pseudomonas koreensis* used as controls to discard competition from non-adapted strains; strains were kindly donated by Dr. Luis Servín and Dr. Gloria Soberón both from *Instituto de Investigaciones Biomédicas*, UNAM. We chose the selecting condition of 16% (w/v) mine-tailing as it reduces 1 fold growth when compared to LB media (Supplementary Figure S3). The selection of plant-derived or complex carbon sources (mannitol) was done by using two independent media culture: 1) Soybean (*Glycine max*) sprouts sterilized homogenate as C,N, and P sole source; 2) LB media with mannitol (3% w/v) as sole carbon source. There were two independent lines of each culture media, each one with duplicates.

The SC was inoculated into 30 ml of each experimental medium. After SC inoculation, the flask was incubated for 48 h (28°C, 70 rpm), then 1 ml of culture was transferred to fresh medium; this procedure was repeated three more times. Then, twenty-two serial passes were performed as described above with the new fresh experimental medium. After the experimental evolution setup, bacterial colonies were re-isolated, and the whole set of isolates was named as the final synthetic community (FSC).

### Metagenomic DNA extraction

Plant roots were vortex-shaked and sonicated on sterile PBS solution, to get the rhizospheric pellet as described elsewhere (Lundberg *et al*.,2012). The metagenomic DNA was extracted from the pellets with the MoBio® PowerSoil extraction kit (MoBio Laboratories, Solana Beach, CA, USA). For the cultured microbes, individual bacterial colonies from each plant, SC, and FSC were tooth-picked into a 2 ml centrifuge tube for each community containing sterilized water, proteinase K and lysozyme (both from Sigma-Aldrich, St. Louis Missouri, United States) were added for an initial lysis (37°C, 30 min). The tubes were centrifuged and the pellets were used as the source for DNA extraction with the MoBio® PowerSoil extraction kit.

### Amplification of the 16S rRNA gene

The PCR reactions were carried out to amplify the V3-V4 region (341F and 805R primers; 464 bp amplicon; Klindworth 2013) following the Illumina® MiSeq™ protocol with 5’ overhangs. We performed triplicated PCR reactions for each sample, using the high fidelity *Pfx platinum* polymerase (Invitrogen, Thermo Fisher Scientific Corporation, Carlsbad, California, USA) with following conditions: denaturation at 95 °C for 3 min; 5 cycles of denaturation at 94 °C for 30 s, annealing at 55 °C for 30 s, and extension at 68 °C for 30 s, 25 cycles of two-step cycling with denaturation at 94 °C for 5 s, and extension at 68 °C for 30s; final extension at 68 °C for 5 min. Finally, amplicons from each replicate were pooled and purified with the Wizard SV Gel and PCR Cleanup System kit (Promega Corporation, Madison, Wisconsin, USA).

### DNA sequencing

Amplicons of the 16S rRNA gene were sequenced through Illumina MiSeq 2×250 paired-end technology (Illumina®, San Diego, California, USA) at the Unidad de Secuenciación Masiva of the UNAM Biotechnology Institute (Cuernavaca, Morelos, Mexico). The whole shotgun metagenomic DNA from the rhizosphere of *A. farnesiana* growing on mine tailing substrate and the SC were sequenced with Illumina® HiSeq 2000 (2×100) at Macrogen (Korea). Finally, the shotgun metagenomic DNA of the FSC community was sequenced at the Genomic Services laboratory *Unidad de Genómica Avanzada* (UGA, formerly LANGEBIO-CINVESTAV, Mexico) using Illumina® MiSeq 2×300 technology.

### Bioinformatic analyses

Quality control of the 16S rRNA gene amplicon sequences was performed with FASTX-toolkit v0.0.14 (http://hannonlab.cshl.edu/fastx_toolkit/) and paired reads were merged with PANDAseq v2.11 (Masella *et al*., 2012). OTUs were built clustering the merged reads at 97% identity with *cd-hit-est* v4.6 (Fu *et al*., 2012). The QIIME v1.9.1 pipeline (Caporaso *et al*., 2010) was used to assign taxonomy using the BLAST v2.2.22 algorithm (Camacho *et al*., 2009) against the Greengenes database (release 13_8; DeSantis *et al*.,2006) and sequences from chloroplasts and mitochondria were removed. The resulting OTUs table was analyzed with the phyloseq library v1.24.2 (McMurdie & Holmes, 2013) for R v3.5.1 (www.r-project.org) removing singleton OTUs.

The whole metagenome shotgun (WMS reads) were processed with Trimmomatic v0.33 (Bolger, Lohse, & Usadel, 2014), keeping sequences with minimum length of 36 nucleotides and a Phred quality > 15 in sliding windows of 4 nucleotides. Whole shotgun metagenomic taxonomic profile was done using high-quality reads (HQ-reads), then mapped them against the NCBI-nr-protein database with the Kaiju v1.6.2 program (Menzel, Ng & Krogh, 2016). Additionally, WMS 16S rRNA gene fragments were recovered from the HQ-reads with SSU-ALIGN v0.1.1 (Nawrocki & Eddy, 2013) and taxonomy was assigned with BLAST v2.2.22 (Camacho *et al*., 2009) against the Greengenes database (release 13_8; DeSantis *et al*.,2006).

HQ-reads of each sample were assembled separately with MEGAHIT v1.1.3 (Li *et al*., 2016), using minimum kmer size = 21, maximum kmer size = 141, kmer increment = 12, and removing unitigs with average kmer depth < 2, and the resulting contigs were used to predict ORFs and protein sequences with Prodigal v2.6.3 (Hyatt *et al*., 2010). Protein sequences were clustered at 90% identity with CD-HIT v4.7 (Fu *et al*., 2012) and the representative sequences were annotated against the M5nr database (Wilke *et al*., 2012) using DIAMOND v0.9.22 (Buchfink, Xie, & Huson, 2014). An independent annotation was also done using the BlastKOALA server (Kanehisa, Sato & Morishima, 2016). Unannotated proteins of all samples were clustered with CD-HIT v4.7 (Fu *et al*., 2012) at 70% identity to add unannotated data to the protein content comparisons. To compare predicted protein content, a protein feature table was built for each metagenome including both the abundance of proteins annotated with the M5nr database and the abundance of unannotated protein families clustered at 70% identity. The abundance of each protein was assigned through mapping of HQ-reads against predicted ORFs with Bowtie 2 v2.3.4.2 (Langmead, & Salzberg, 2012). The gene table was inspected using phyloseq v1.24.2 (McMurdie & Holmes, 2013) for R v3.5.1 (www.r-project.org) through the Subsystems ontology (Overbeek *et al*., 2005) downloaded from the MG-RAST API (Wilke *et al*., 2015).

Metagenomic HQ-reads of the FSC dataset were mapped against the *Enterobacter* sp. SA187 genome (RefSeq accession GCF_001888805.2; Andrés-Barrao, *et al*.,2017) with Bowtie 2 v2.3.4.2 (Langmead, & Salzberg, 2012) and read coverage along the reference genome were visualized with CIRCUS v2.7 (Naquin, *et al*.,2014) and the CGView server (Grant & Stothard, 2008). To recover genomic sequences, metagenomic contigs of all samples were classified with Kraken v0.10.6 (Wood & Salzberg, 2014) and assigned contigs from each analyzed genus was mapped to a reference genome downloaded from the NCBI Assembly database using scaffold_builder (Silva *et al*.,2013), Quast v5.0.0 (Gurevich, *et al*.,2013), and Mummer v3.0 (Kurtz, *et al*.,2004). Detailed bioinformatic and statistical protocols are available at: https://github.com/genomica-fciencias-unam/nacozari/

## Results

### Selection of a Synthetic Community

A total of 861 isolates were recovered from the rhizospheres of the pioneer plants growing on the tailing dam at Nacozari de García. The colony morphology classification resulted in 105 unique morphotypes. Through a multivariate morphology analysis (Supplementary Figure 2), the overall colony morphological diversity was evaluated and a subset of 235 isolates was selected to produce up the Synthetic Community (SC). A microcosm experiment was performed, 22 serial passes were carried out in media supplemented with 16% (w/v) mine-tailings, 144 isolates representing 10 morphotypes were recovered from the Final Synthetic Community (FSC) (Figure 1).

### Sequencing of mine tailing and cultured microbial communities

Environmental rhizosphere microbiomes of plants growing on mine tailings were inspected through the massive sequencing of the 16S rRNA gene to measure the overall in diversity and find out the fraction recovered with our culture-dependent approach. The 16S rRNA gene amplicon sequencing resulted in 2,072,065 raw paired reads, of which 47.76% were assembled and clustered into 21,851 OTUs (Supplementary Table S1). The mine tailing rhizosphere with the largest number of OTUs was the roots from Bc with 8,868, followed closely by Bs with 8,742, while the sample with the lowest number was AfP1 with 3,581. According to the Chao1 theoretical estimation of the maximum number of OTUs, the percentage of recovered OTUs ranged between 85.60 (for AfP1) and 95.86 (for AfP2). Similarly, 9,271 filtered OTUs were found in all cultivated community samples ranging from 2,716 for Bc-CCT to 718 for Gl-CCT. The coverage of observed OTUs was highest for Gl-CCT with 97.81% and lowest for Bc-CCT with 86.85%.

Three samples were analyzed through whole metagenome shotgun sequencing: the mine tailing rhizosphere of *A. farnesiana* growing on the vegetation patch 1 (Af-MG), the SC, and the FSC. The environmental metagenome Af-MG included 33,354,342 reads spanning 3,368,788,542 bp and the GC% content was 58.71%. The SC metagenome comprised 36,478,480 reads adding up to 3,684,326,480 bp with a GC% of 62.02% while the FSC had 20,578,692 reads spanning 6,173,607,600 bp with 59.5 GC% (Supplementary Table 1).

### Changes in diversity from environmental to cultivated microbiomes

In most samples from mine tailing substrate, the dominant bacterial phylum was *Actinobacteria* (46.74%, 39.98% and 38.22% for AfP1, Bc, and Bs, respectively) and showed fewer sequences for *Proteobacteria*, that was the most abundant in AfP2 and Gl (63.78%, and 37.25%, respectively; Supplementary Table S2). The cultured consortia were all dominated by *Proteobacteria* (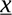=¿80.06%) and *Actinobacteria* to a lesser extent (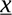=¿18.47%; Supplementary Table 3), this loss of diversity was quantified through the Shannon and Simpson indexes of these samples. Both Shannon and Simpson indexes decayed when compared to their uncultured counterparts, for example, the rhizospheric community of *B. sarothroides* had a Shannon diversity index (H) = 7.14, while the cultivated community had H = 4.11 (Figure 2).

**Figure 2.**
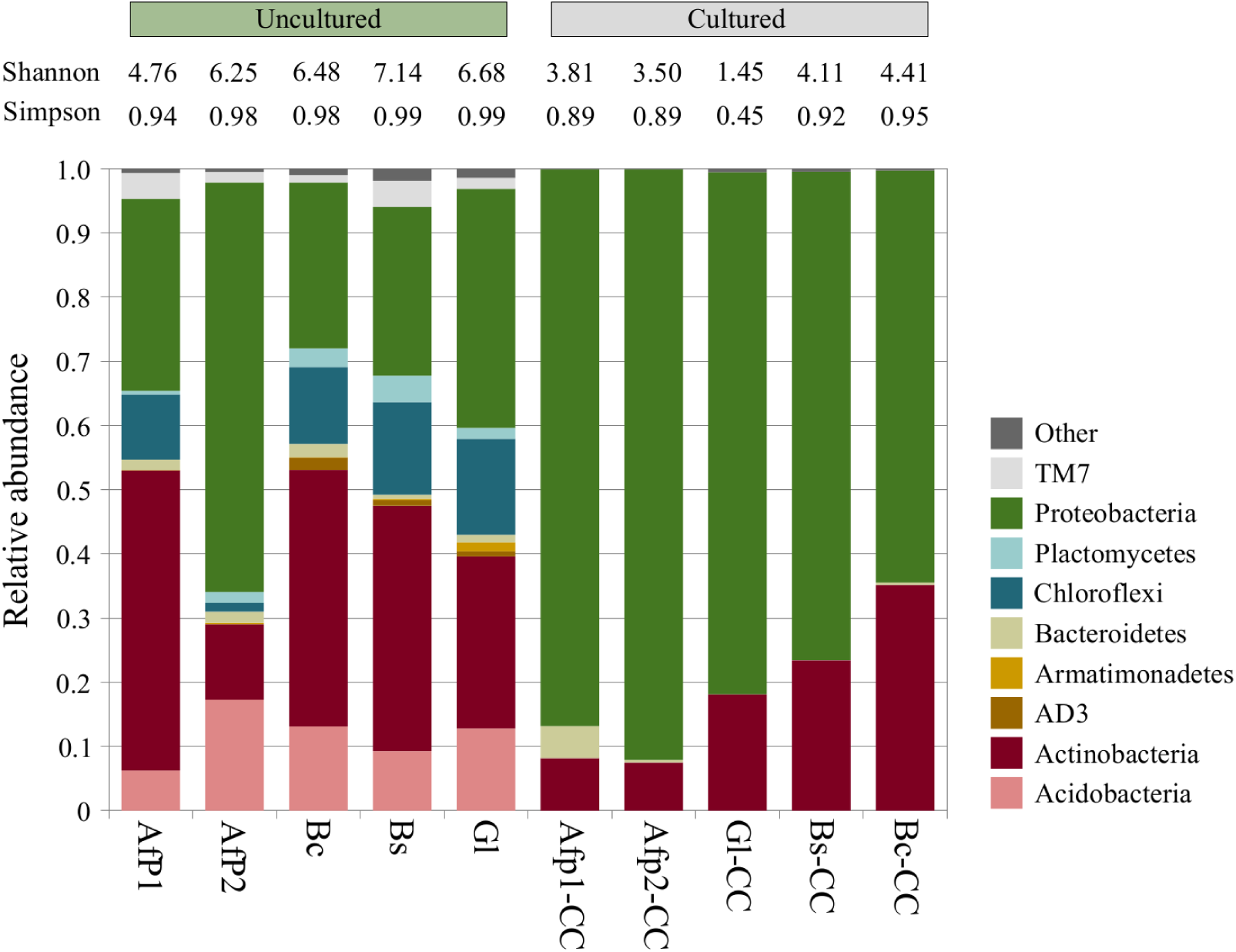
Taxonomic diversity is substantially decreased in the cultured communities, which are dominated by Proteobacteria. Relative frequency of the most abundant phylotypes for 16S rRNA gene amplicon sequences. Values for Shannon and Inverse Simpson’s diversity indexes for OTU level counts are shown. Af: *A. farnesiana*; Bc: *B. coulteri*; Bs: *B. sarothroides*; Gl: *G. leucocephalum*; -CC: cultivated community.

A total of 94 genera were identified in the SC using only the V3-V4 16S rRNA sequences, sharing 18 genera out of 43 in FSC (Figure 3B, Supplementary Table S4), from which the most frequent were also found in all the plants growing in the tailing vegetation patch, such as *Pseudomonas* and *Ralstonia* (17.39 and 9.69%, respectively). Other abundant genera, like *Enterobacter*, were only observed in the rhizosphere of some plants such as *B. coulteri*. The SC also included 10 unique genera, which were all observed at a percentage < 0.03% of the WMS-reads. Additionally, most HQ-reads (54.34 %) of the FSC were mapped to *Enterobacter* sequences (Figure 3C; Supplementary Table 5).

**Figure 3.**
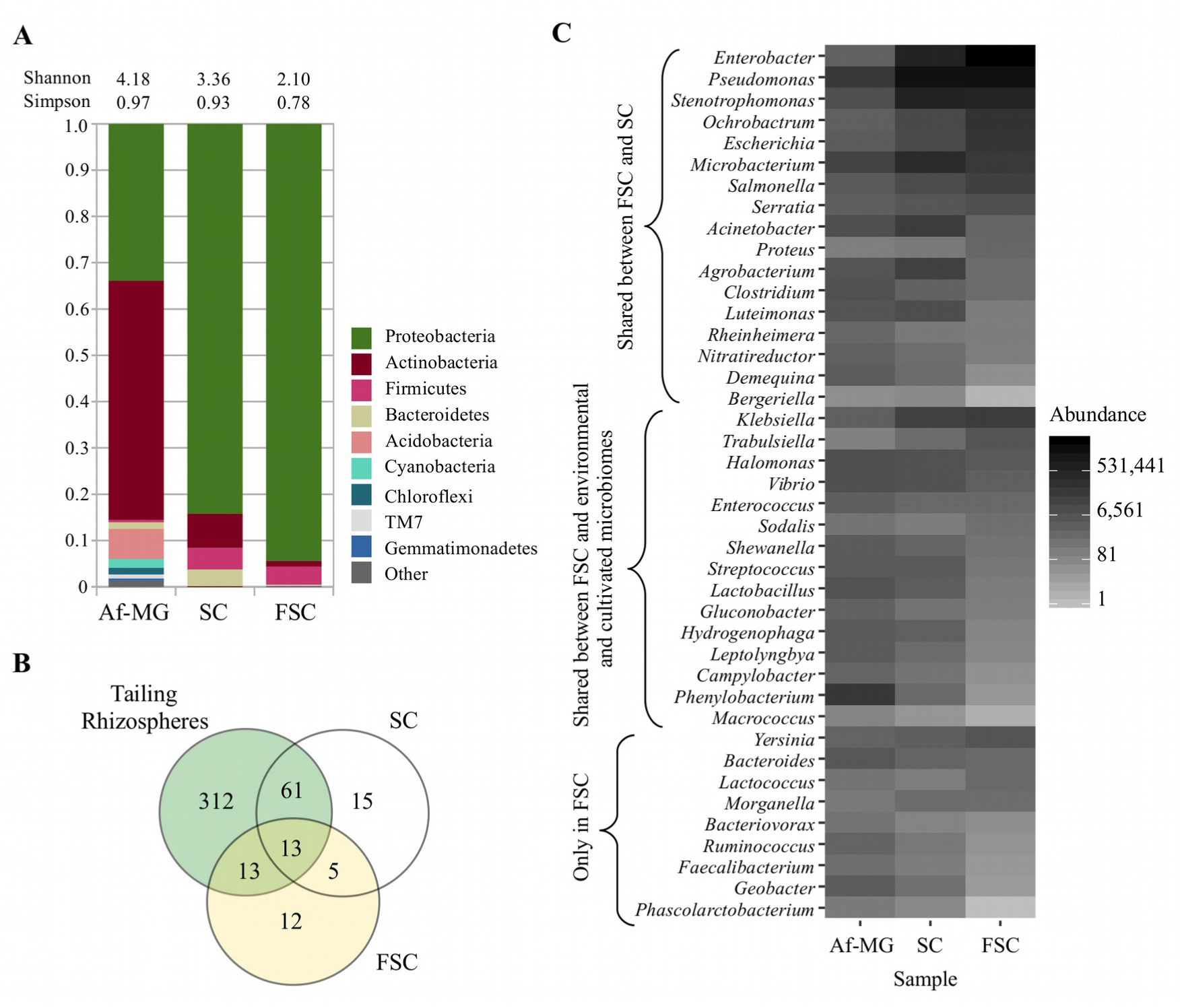
Taxonomic diversity is substantially decreased in the FSC and is dominated by proteobacterial taxa. (A) Relative frequency of bacterial phylotypes of binned 16S rRNA gene (V3-V4) shotgun metagenome sequences from wild samples (Af-MG) and the synthetic communities. (B) Shared genera from 16S rRNA gene assignments, from the 4 rhizospheres of local plants (tailing rhizospheres amplicons), and shotgun fragments binned to V3-V4 region from the synthetic communities. (C) Metagenomic binning and taxonomic placement using translated genes searched in the NCBI NR protein database for the 41 found genera reported for the final synthetic community (FSC).

Nonbacterial sequences in the metagenomic samples were also found: eukaryotic reads represented the 2.39% in the Af-MG, 0.14% in the SC and 0.06% in the FSC samples. The genera with the highest number of mapped metagenomic reads in all three metagenomes were fungi: *Rhizophagus* with 14,965 reads in Af-MG (with 54 and 4 reads in the SC and FSC metagenomes, respectively); *Coniochaeta* with 2,600 reads in SC (567 and 2 reads in Af-MG and FSC samples, respectively); and *Rhizoctonia* with 613 reads in FSC (2,641 and 525 reads in Af-MG and SC datasets, respectively). Sequences from the Archaea domain were also found representing 0.24%, 0.02% and 0.005% of the metagenomic reads in the Af-MG, SC, and FSC samples, respectively. In all samples, the most abundant group was the phylum *Euryarchaeota* and *Halobacteria* class (Supplementary Table S5).

### Metagenomic profiles differences from environmental to selected cultivated communities

Predicted protein diversity of the metagenomes was compared through proteins finding homologues in the reference M5nr database, including hypothetical and conserved hypothetical proteins. Additionally we incorporate, the protein sequences clustering (70% amino acid identity) of predicted hypothetical proteins without homologue protein matches in the M5nr. The protein diversity was described by the number of observed and shared proteins between wild and synthetic communities, along the protein Shannon diversity index (Figure 4A). A total of 318,157 proteins were found: 225,733 annotated proteins with the M5nr database and 92,424 hypothetical protein clusters. The environmental metagenome Af-MG had the highest count of proteins, followed by the SC (with 173,717 and 146,928 proteins, respectively). Interestingly, the SC had a higher Shannon index than the Af-MG metagenome (11.04 and 11.03, respectively). Even though, the FSC presented the lowest observed proteins (23,196) and Shannon index (8.78), it matched the most proteins to the M5nr database (90.78%), compared with SC (77%), and the Af-MG (63.35%).

**Figure 4.**
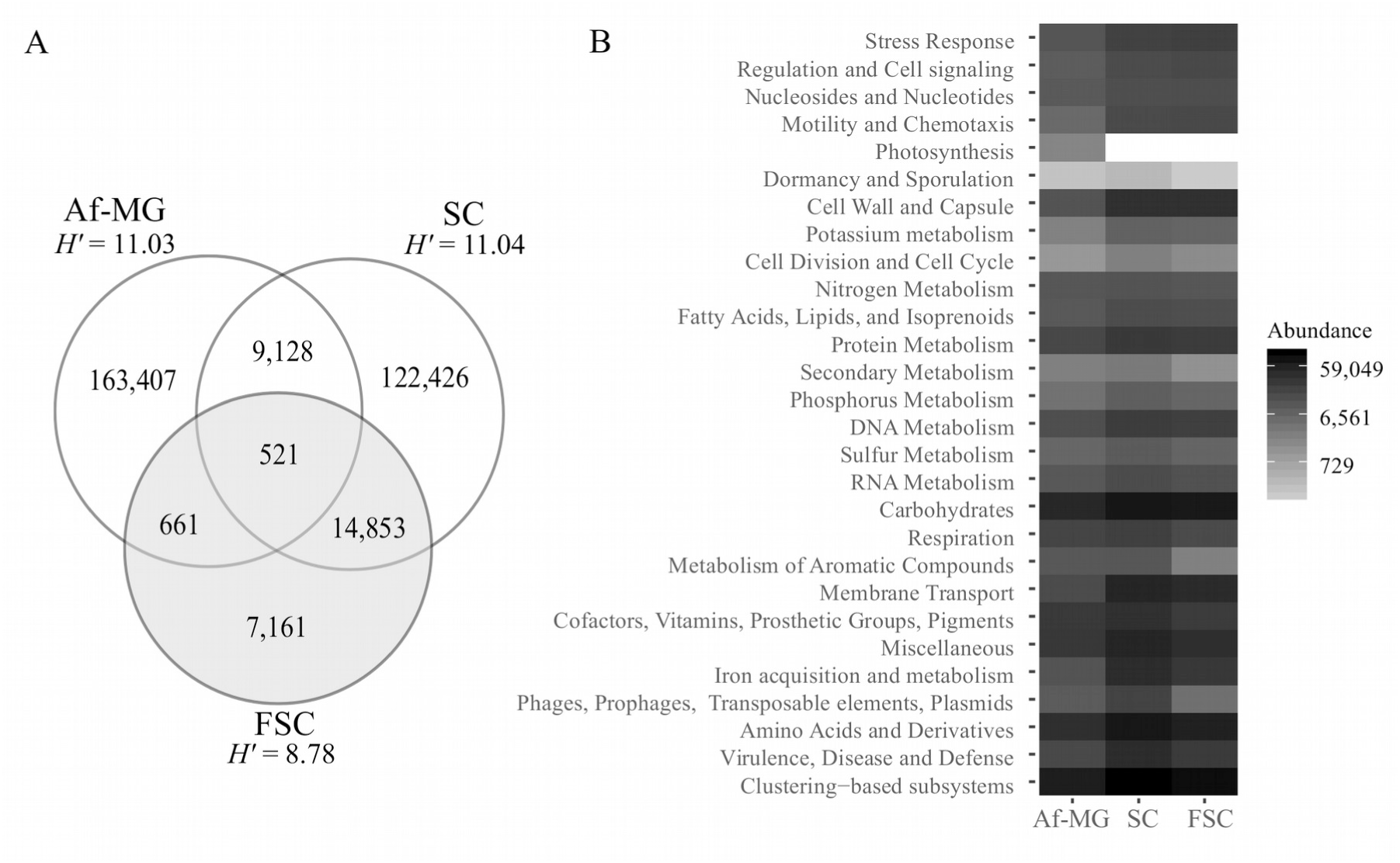
The selected synthetic community (FSC) reduces wild and initial protein diversity and is enriched on subsystems associated with stress response, motility and lacks photosynthesis genes. (A) Total predicted, shared, and unique proteins for wild *Acacia farnesiana* rhizosphere metagenome (Af-Mg), compared to the protein numbers of the culturable SC and FSC communities. Shannon’s protein diversity index (*H’*) is decreased in the FSC compared to Af-Mg and SC. (B) Metagenomic functional classification showing thousands of reads mapped to genes associated with the first level of SEED Subsystems.

A core set of 521 proteins were shared between all metagenomes, of which 166 were annotated by homology with the SEED database, including arsenic resistance proteins, metallic cation efflux systems, antibiotic resistance factors like the polymyxin resistance protein ArnT, and carbohydrate ABC transporter proteins (Supplementary Figure S5, Supplementary Table S6). Among these proteins, a probable Co/Zn/Cd efflux system membrane fusion protein was found enriched in the FSC with 5.95% of mapped reads in comparison with the 3.66% of reads in the SC metagenome.

Classification with the SEED subsystems ontology revealed enriched categories for the synthetic communities such as membrane transport, amino acid and derivatives, cell wall and capsule, and Iron acquisition and metabolism (Figure 4B). In the stress response, and motility and chemotaxis subsystems, the FSC was the sample with the highest normalized read counts, with 3.68 % and 2.65 % of mapped reads, respectively. Specifically, from the stress response subsystem, the osmoprotectant ABC transporter YehZYXW (Lang *et al*.,2015) was most abundant in the FSC metagenome with 0.2% of mapped reads, while the SC mapped 0.02% and was absent in the Af-MG metagenome. From the amino acid and derivatives subsystem, proteins involved in amino acid degradation of arginine and ornithine were enriched in the FSC with 1% mapped reads (against 0.84 and 0.5% of mapped reads in the SC and Af-MG, respectively). The FSC was also enriched in the conjugative transfer (1.34% of mapped reads) and ABC di-peptide transporters (0.87%), from the membrane transport subsystem, and for enterobactin biosynthesis (0.58%). The FSC had a high frequency of functions from the miscellaneous subsystem (6.39% of mapped HQ-reads), including the ACC deaminase (COG2515), as well on the protein-export related proteins from the Clustering-based subsystems. Besides the enriched subsystems, some categories were only observed in the environmental Af-MG metagenome, such as the nitrogen fixation subsystem (including NifH, NifE, NifN, NifW and NifO; Supplementary Figure S6), and other categories were contained in the SC but were lost after the mesocosm experiment, like the coenzyme F420 synthesis subsystem. Similarly, the Indoleacetamide hydrolase (involved in auxin synthesis) was found in the Af-MG and SC metagenomes but was lost after the mesocosm experiment (Supplementary Figure S6).

### Genome reconstruction of *Enterobacter sp*. Nacozari

Since most (54.34 %) of the mapped metagenomic reads of the FSC mapped against sequences of the *Enterobacter* sp. and most (37.14 %) of them aligned to *Enterobacter* sp., SA187 (RefSeq accession GCF_001888805.2; Andrés-Barrao *et al*., 2017), we used this genome as reference to perform comparative genomics. The new strain was named *Enterobacter* sp. Nacozari. A total of 93.78 % of the nucleotides of the reference SA187 genome were represented in 7,356,403 of our metagenomic reads (Figure 5A) with a mean sequence identity of 99.1 %. Mapped loci included 90.71 % of the coding sequences in the reference SA187. The coding sequences for several plant symbiosis proteins reported in the SA187 genome were mapped, including CsgBAC and CsgDEFG (curli fiber subunits and secretion proteins, respectively; Figure 5B), AcrAB (multidrug efflux pump; Figure 5C) and EntCEBAH (enterobactin biosynthesis; Figure 5D) as well as the siderophore exporter protein EntS. Other plant-symbiosis related proteins such as UbiC (chorismate pyruvate-lyase), along with oxidative stress response proteins like SOD1/2 and KatE were included in our sequencing data. Absent loci of the chromosome of *Enterobacter* sp. SA187 in our metagenomic reads included coding sequences from prophage loci such as the phage tail protein, the phage baseplate assembly protein V, and the phage repressor protein CI (Supplementary Figure S7). Additionally, the coding loci of the SA187 strain, DndEDCB (DNA sulfur modification) proteins remained unmapped (Supplementary Figure S7).

**Figure 5.**
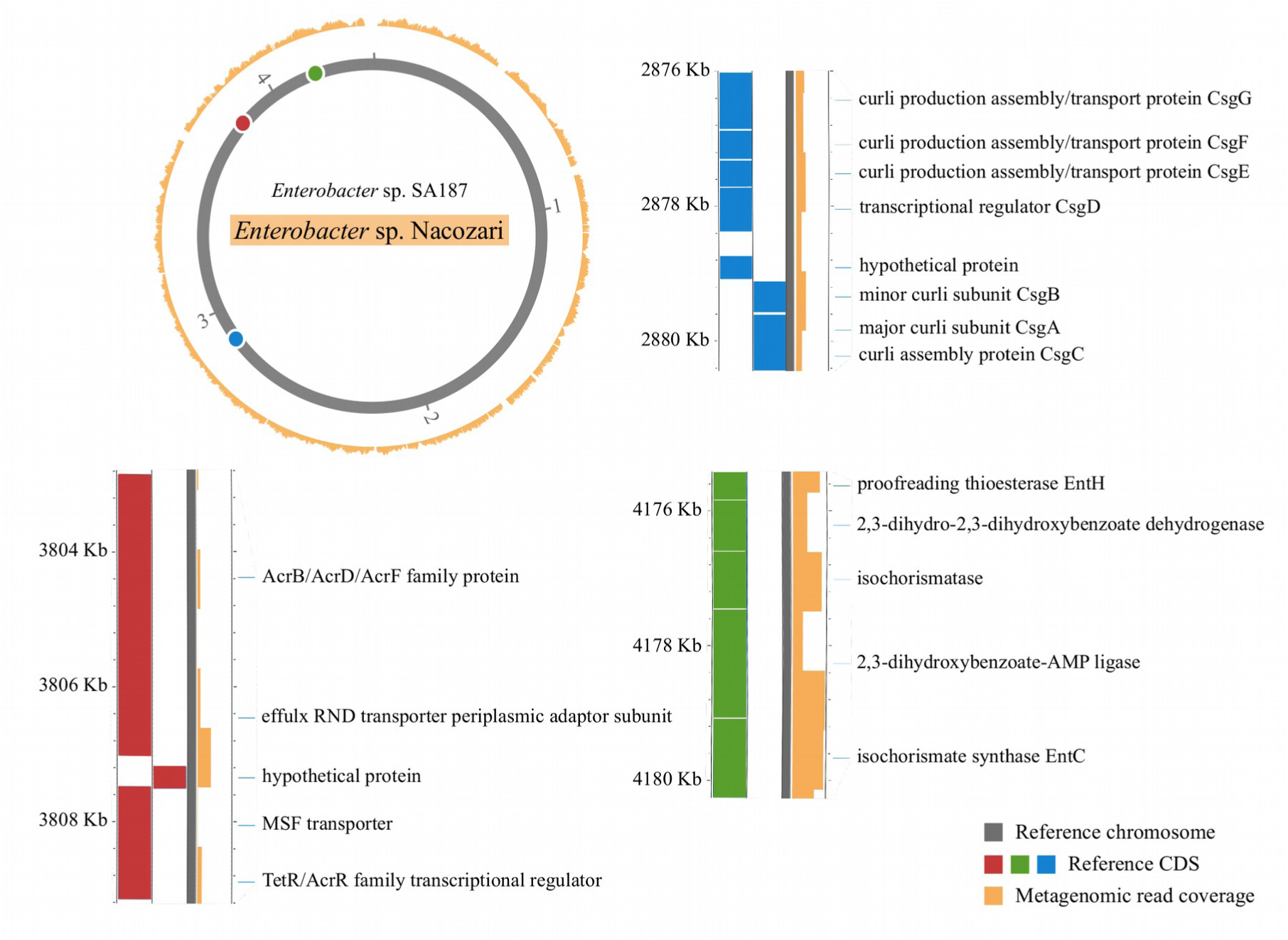
Metagenomic recruitment and assembly of *Enterobacter* sp. Nacozari against *Enterobacter* sp. SA187 reference genome, an already described plant-growth promoter bacteria, including described genes associated with plant growth promotion. *Enterobacter* sp. Nacozari metagenomic sequences (outer yellow ring) aligned to the *Enterobacter* sp. SA187 genome as a reference (scale in Megabases). Selected loci coverage involved in plan-growth promotion in *E*. sp. SA187 are shown like curli fiber subunits (*csgBAC)* and secretion genes (*csgDEFG)*; the multidrug efflux pump *acrAB*; and the enterobactin biosynthesis *entCEBAH*.

To recover additional genes that might be present in our assembly but absent in the reference *Enterobacter* sp., SA187 metagenomic contigs of the three metagenomic samples were classified to get all the *Enterobacter* assigned sequences. The 344 contigs that were tagged as belonging to the *Enterobacter* genus represented 80.14% of the reference genome while the five built scaffolds covered 48.33% of the reference genome (Supplementary Figure S8; Supplementary Table S7) and had a G-C content of 56.38%. A total of 5,150 proteins were predicted from these scaffolds and annotation against the KEGG GENES database revealed that the coding sequences belonging to PqqBCDE, pyrroloquinoline quinone biosynthesis proteins were found within this genome fragments (Supplementary Figure S9; Supplementary Table S8).

## Discussion

We are reporting the Nacozari mine tailing rhizospheric culturable and culture-free taxonomic and metabolic diversity. From the culturable organisms, we selected a synthetic microbial community capable of growing on mine tailing substrate, with plant-derived C, N, and P sources. While rhizospheric communities had high frequencies of Actinobacteria (37.99% on average), as has been observed in our Af-Mg, and arid soils (Crits-Christoph *et al*., 2013), the SC was dominated by *Proteobacteria*. Despite other studies have also recovered isolates belonging to *Enterobacter, Pseudomonas*, and *Chryseobacterium* from heavy-metal contaminated microenvironments (Toribio-Jiménez *et al*., 2014; Matlakowska & Sklodowska, 2009). We also isolated bacteria of plant-associated genera like *Ralstonia* and *Burkholderia* in high frequency, some *Ralstonia* species are well-known pathogens of plants (Hayward, 1991), although no canonical pathogenesis factors such as Egl, PehA/B or CbhA were found in our shotgun metagenomic samples, and *Burkholderia* had been reported as a diazotrophic symbiont (Gillis *et al*., 1995). *Pseudomonas putida*, was the reference genome with the most mapped reads in the SC metagenome, it has been found in soil and rhizospheric communities and recent studies comparing several strains have concluded that this bacterial taxon’s core-genome lacks virulence factors and can catabolize complex carbon sources, including aromatic compounds (Udaondo *et al*., 2016). Similarly, in the FSC, most metagenomic reads were mapped to the genome of *Enterobacter* sp. SA187. This bacterial strain was isolated from the root endosphere of a desert plant and showed plant growth promoting activity (Andrés-Barrao *et al*., 2017). Furthermore, its genome sequence revealed proteins associated with oxidative stress tolerance, antibiotic production, plant hormone regulation, and adhesion (Andrés-Barrao *et al*., 2017).

Even though archaeal sequences were observed at low frequencies (0.24%, 0.02% and 0.005% of the mapped metagenomic reads in the Af-MG, SC, and FSC samples, respectively), organisms of this kingdom have been observed to be tolerant to heavy metals and might be essential helping other organisms to grow in heavy metal contaminated environments (Li *et al*.,2017). Fungal organisms were represented in the metagenomic samples (1.59 %, 0.17 %, and 0.04 % of mapped reads in the Af-MG, SC, and FSC samples, respectively) including the *Rhizophagus* genus, which is a well-known arbuscular mycorrhizal symbiont of plants that has been shown to stimulate plant growth in *Acacia holosericea* (Duponnois *et al*., 2005), that was the fungi with most mapped sequences in the Af-MG metagenome but its frequency decreased drastically in the cultured samples with 14,965 reads in Af-MG (0.09 %), 54 in the SC and 4 reads in the FSC metagenome.

We used the Shannon diversity index to describe metabolic diversity, analyzed through matched proteins against the M5nr database and the clustering of unmatched proteins at 70% sequence identity. Although the ranges of the Shannon index varied, the same pattern was observed: the Af-MG and the SC had similar values while the FSC had a lower value (11.03, 11.04 and 8.78 for Af-MG, SC and FSC, respectively). The metabolic diversity in the SC metagenome, resembling the value of the culture-free Af-MG, may be due to the fact that SC resulted from the combination of several cultured rhizospheric communities, from four plant species and that each one carried a specific functional profile which added up to the unexpectedly high number of unique protein features in this sample (Figure 4A). Interestingly, the taxonomic diversity in the SC was lower (3.36) than the one in the Af-MG sample (4.18). This result is in opposition to previous observations where the functional diversity strongly correlates with the taxonomic diversity (Fierer *et al*.,2013), and might be a result of similar taxa carrying different genes (*i*.*e*. a large pan-genome), as has been observed for the *Pseudomonas* genus (Hesse *et al*., 2018), which was present in high frequency in the SC metagenome (17.39 % of mapped reads).

A total of 521 proteins were present in the Af-MG, SC, and FSC, thus being a shared core between environmental metagenomes and cultured ones. The core features proteins that have been reported as heavy metal resistance factors such as the CusA P-type ATPase (Taylor *et al*.,1988; Gillan *et al*.,2015) and the As resistance protein ArsH (Chen, Bhattacharjee & Rosen, 2015; Li *et al*.,2014). Furthermore, within this core we also found antibiotic resistance genes such as beta-lactamases and the polymyxin resistance protein ArnT which might be essential to withstand the antagonism in these microbial communities, they could also be co-selected along with the heavy metal resistance genes (Pal *et al*., 2015).

The stress response proteins in the FSC included stress response factors which were selected in the harsh conditions of the mesocosm experiment. Among them, we found YehZ, an osmoprotectant transporter found in *Enterobacteriaceae* that has been upregulated in nutrient starvation, acidic pH, and hyperosmotic stress conditions (Kim, Ryu & Yoon, 2013). Nutrient acquisition strategies which may be relevant in a plant-host associated environment, were also selected in this system. Such as TonB-dependent receptors which are involved in the uptake of dissolved organic matter, siderophores and vitamins in marine environments (Tang *et al*.,2012). Likewise, peptide ABC transporters have also been found enriched in genomes from rhizospheric bacteria (Matilla *et al*.,2007), and bacterial secretion systems which might be relevant in the bacterial interactions with the plant host and other bacteria (Green & Mecsas, 2016). Other proteins associated with plant-bacteria symbiosis found in the FSC are siderophore producing enzymes (Kloepper *et al*.,1980), and the ACC deaminase, which modulates ethylene levels in the plant, thus preventing the negative effects of this plant hormone in stressing conditions (Mayak, Tirosh & Glick, 2004). This community also harbors galactoglucan biosynthesis enzymes, that are involved in biofilm formation, which might form complexes with the metal ions from the mine tailing substrate and thus preventing their accumulation by the plant (Macaskie & Dean, 1987). However, other crucial plant growth promoting proteins were lost after the synthetic ecology experiment, like the indoleacetamide hydrolase, involved in the synthesis of the plant hormone indole-3-acetic acid (Clark *et al*., 1993).

The strain *Enterobacter* sp. SA187 has been described as an endophytic plant growth promoting organism which could stand unfavorable abiotic conditions such as oxidative stress (Andrés-Barrao *et al*., 2017). Our metagenomic reads mapped to plant symbiosis proteins such as AcrAB, which form an efflux pump involved in plant colonization (Burse, Weingart & Ullrich *et al*., 2004); and CsgBAC, that code for the curli fiber subunits that are required during late plant root colonization and might mediate the adhesion of this organism to the plant surface (Cowles *et al*.,2016). The enterobactin biosynthesis polycistron *entCEBAH*, recovered from the metagenomic reads of the FSC, have been shown to promote plant growth in heavy metal contaminated soil through the provision of Fe in the presence of other metals and through the decrease of oxidative stress in the plant due to binding of metals near the roots (Dimkpa *et al*.,2009). Thus, *entCEBAH* might be a consistent plant growth promoting factor in mine tailings as siderophores. Besides, the enterobactin exporter EntS has been reported as highly expressed when *Enterobacter* sp. SA187 was associated with roots of *Arabidopsis thaliana* (Andrés-Barrao *et al*., 2017). Indirect plant growth promoting proteins were also included, like UbiC, which is involved in the synthesis of 4-hydroxybenzoate, an antibiotic that has been shown to reduce the rate of infection of *ubiC*-transformed potato plants of the fungal pathogen *Phytophthora infestans* (Köhle *et al*.,2003). Additionally, proteins relevant to the mine tailing conditions were also present in these predicted protein sequences, including SOD1/2 and KatE; however, these proteins could also be relevant in plant tissue establishment, as a plant defense mechanism is the production of reactive oxygen species (Keppler, Baker & Atkinson, 1989).

The metagenomic reconstructed genome, *Enterobacter* sp. Nacozari, has the metabolic potential to assimilate plant-derived carbohydrates such as mannose, fructose, and starch. A diverse array of monosaccharide and amino acid ABC transporters was also observed, which could be mediating the capture of plant rhizodeposits and have been found enriched in a plant associated *P. putida* (Matilla *et al*.,2007). Moreover, we identified several proteins that have been previously described as plant-growth promoting factors such as the PqqBCDE proteins, that contribute to mineral phosphate solubilization in *Enterobacter intermedium* (Kim *et al*.,2003). Furthermore, our following work is to test the plant-growth promoting activities of single isolates as well as the whole FSC under greenhouse experiments with plants occurring in the Nacozari mine tailings.

## Conclusion

The taxonomic profile of the mine tailing rhizosphere communities shifted from the environmental dominance of *Actinobacteria* to *Proteobacteria* in the cultivated consortia, while a decreasing diversity gradient was observed from the environmental microbiomes to the FSC. In general, the metabolic potential of the selected microbial community (FSC) was enriched in genes related to membrane transport, amino acid metabolism, and iron acquisition in comparison with the Af-MG environmental metagenome and was enriched in motility and stress response subsystems in comparison with the initial synthetic community. A core of genes shared by the environmental and cultivated metagenomes included heavy metal homeostasis related proteins and antibiotic resistance enzymes that could be the result of selective constraints within the mine-tailing. The FSC harbored functions related to plant-growth promotion such as siderophore production and genes coding for plant hormone synthesis. Finally, we assembled the metagenome-derived genome of the *Enterobacter* sp. Nacozari, in which we found direct and indirect plant-growth promotion predicted coding genes. The metabolic potential of the FSC presents promising features that might make it useful for plant-growth promotion in a phytostabilization strategies. Further work is in progress to evaluate the plant-growth promotion of the FSC.

## Data availability

The whole metagenomic reads area available in the DDBJ/EMBL/Genbank databases under Bioproject accession: PRJNA525709; BioSample accessions: SAMN11174940, SAMN11174941, SAMN11174942, SAMN11174943, SAMN11174944, SAMN11174945, SAMN11174946, SAMN11174947, SAMN11174948, SAMN11174949, SAMN11174950, SAMN11174951, and SAMN11174952. Detailed bioinformatic and statistical protocols are available at: https://github.com/genomica-fciencias-unam/nacozari/

## Acknowledgments

We thank Dr. Julio Campo at *Instituto de Ecología*, Universidad Nacional Autónoma de México (IE-UNAM) for his assistance processing mine tailing material. Rodrigo García Herrera, head of the Scientific Computing Department at LANCIS-IE-UNAM, for running the HTC infrastructure used for the analyses. MR is a student from the Posgrado en Ciencias Biológicas, Universidad Nacional Autónoma de México (UNAM) and received fellowship 692969 from CONACyT.

## Author Contributions

LDA, DG, MP, RCO, and FMF conceived and designed the experiments. Field sampling, and laboratory work was coordinated and executed by: DG, MR, JB, AL, JMF, HB, CHK, RCO, FMF, LDA. DG carried out the microbiology/mesocosm experiment. Bioinformatic and statistical analysis was performed by MR, DG, JB, AL, LDA. Contributed reagents/materials/analysis tools: FMF, MP, RCO, LDA. MR and LDA were responsible for primary manuscript elaboration and merging input from all the authors. MR, DG, MP, and LDA contributed to figure elaboration.

