## Supplementary Figures for "Rhizosphere metagenomics of mine tailings colonizing plants: assembling and selecting synthetic bacterial communities to enhance *in situ* bioremediation"

A

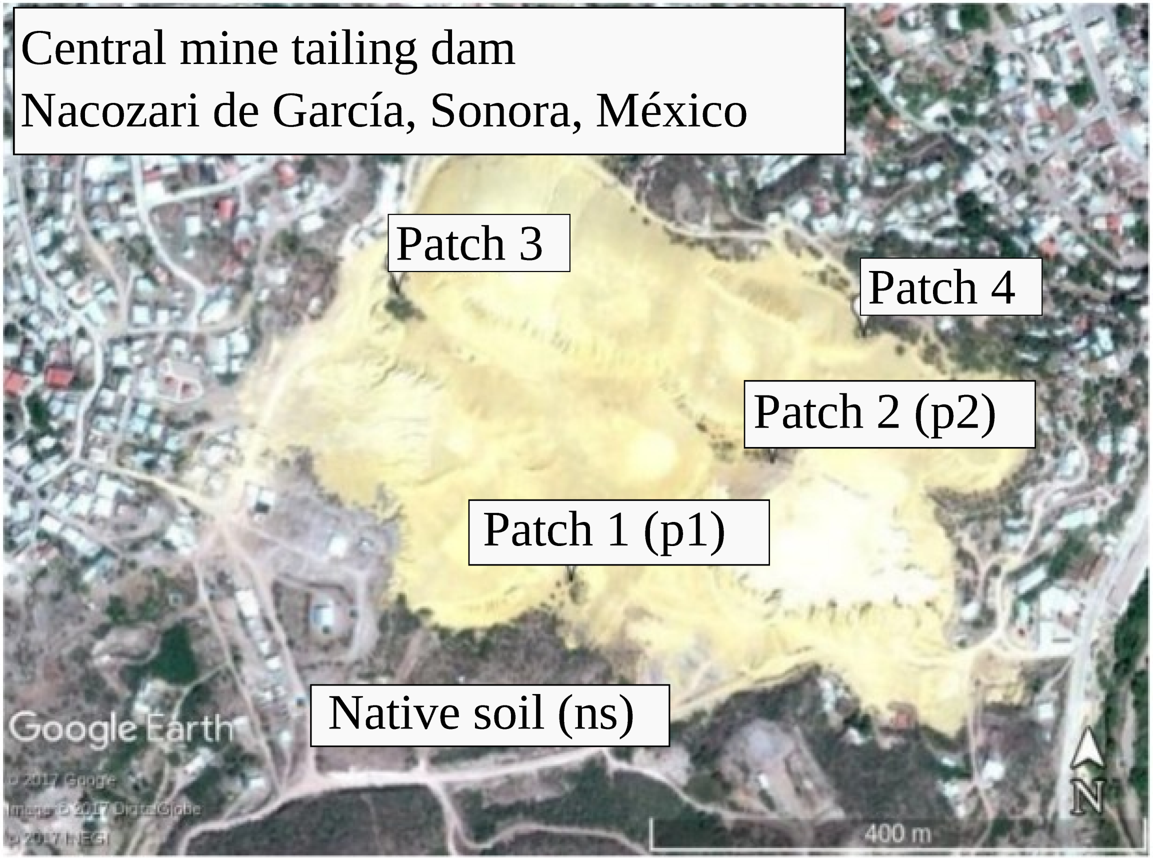


B

| Plant species, sampling location | Whole metagenome shotgun (MG) | V3-V4 16S rRNA gene amplicons | Cultivated community (CC) |
|---|---|---|---|
| *Acacia farnesiana*, p2 (Afp2) | | ✓ | ✓ |
| *Acacia farnesiana*, p1 (Afp1) | ✓ | ✓ | ✓ |
| *Brickellia coulteri*, p1 (Bc) | | ✓ | ✓ |
| *Baccharis sarothroides*, p1 (Bs) | | ✓ | ✓ |
| *Gnaphalium leucocephalum*, p1 (Gl) | | ✓ | ✓ |
| *Acacia farnesiana*, ns (Af-ns) | | ✓ | ✓ |
| Synthetic Community (SC) | ✓ | | |
| Final Synthetic Community (FSC) | ✓ | | |

**Supplementary Figure S1.** Sampling location and overview. (A) Sampling location. (B) Processed samples

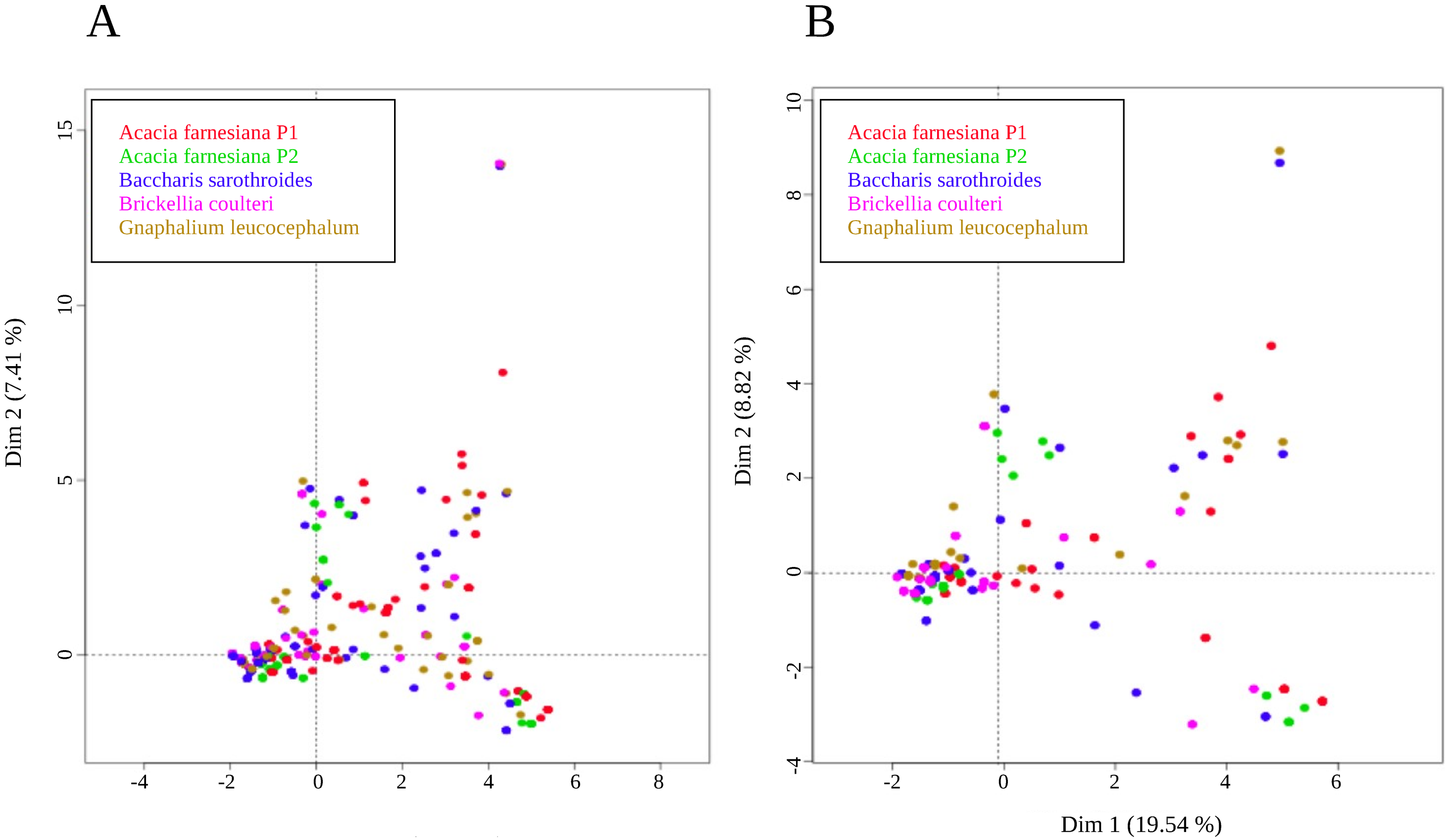


**Supplementary Figure S2.** Comparison of the colony morphology diversity between the original cultivated consortia and the SC. Multiple classification analysis of the morphological data of the (A) 861 isolates of the cultivated consortia of the 5 plants and (B) the 235 isolates that formed the SC

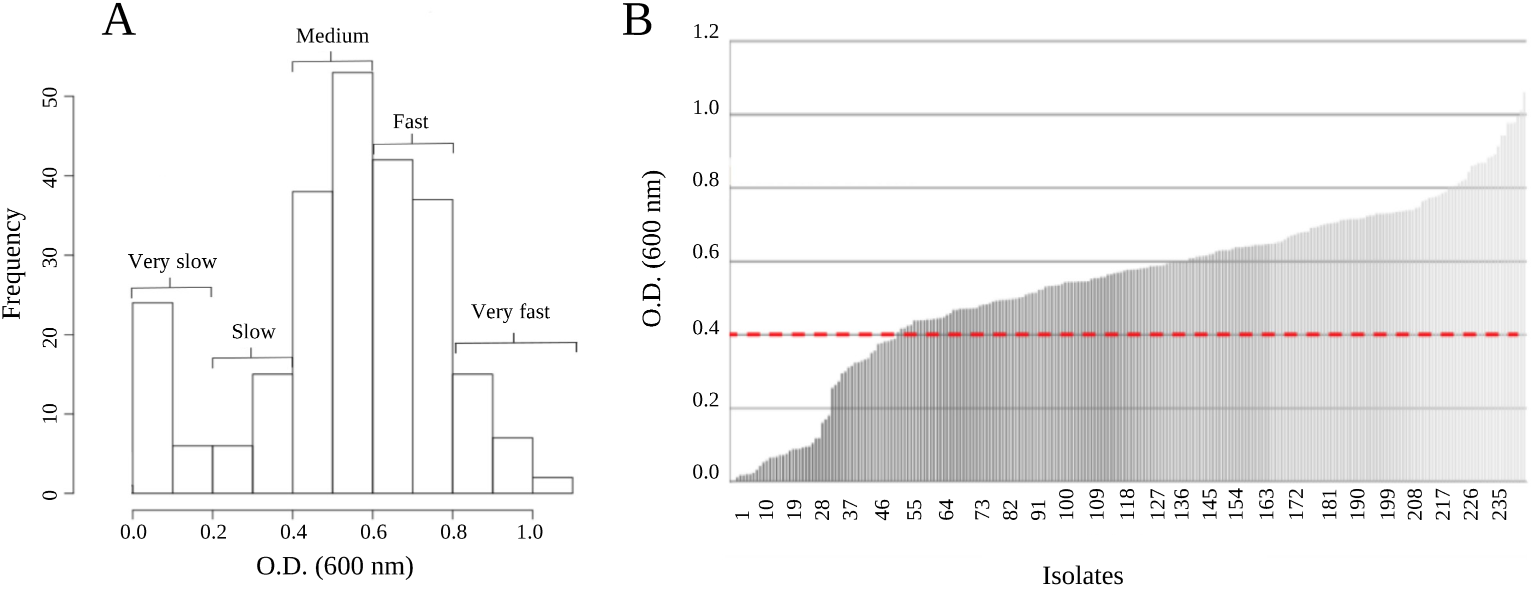


**Supplementary Figure S3.** Growth rates of the 235 members of the SC. (A) Histogram of the O.D. (λ 600 nm) of the 235 colonies of the SC after 18 h. The definition of growth rates was as follows: very slow 0 < O.D. < 0.2; slow 0.2 < O.D. < 0.4; medium 0.4 < O.D. < 0.6; fast 0.6 < O.D. < 0.8; very fast 0.8 < O.D. < 1.0. (B) Maximal O.D. after 42 h.

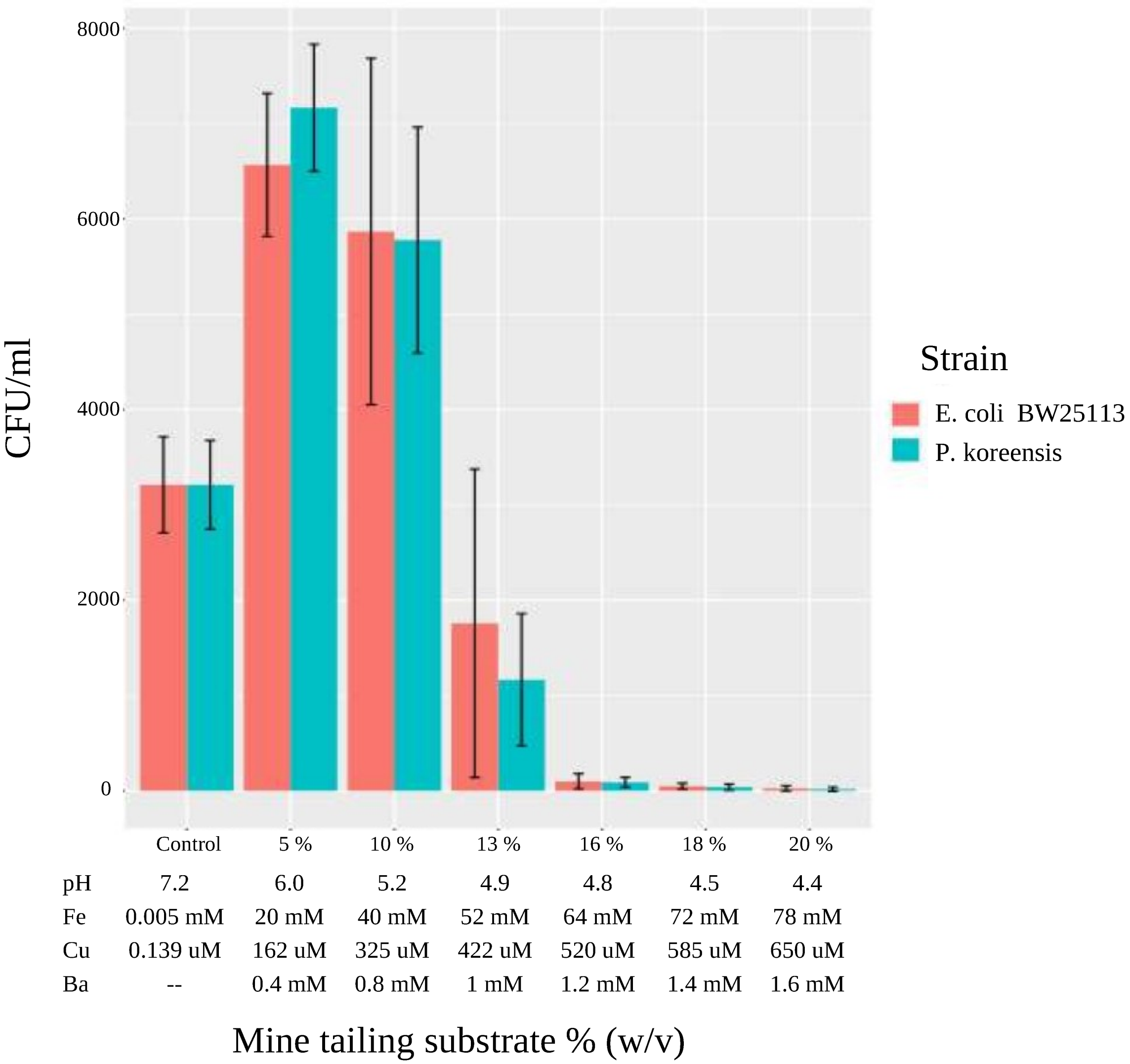


**Supplementary Figure S4.** Effect of mine tailing substrate at different concentrations on the growth of *Escherichia coli* BW25113 and *Pseudomonas koreensis*. Counted CFU/ml after a growth assay on control medium (without mine tailing substrate) and media with 5, 10, 13, 16, 18, and 20% w/v of mine tailing substrate.

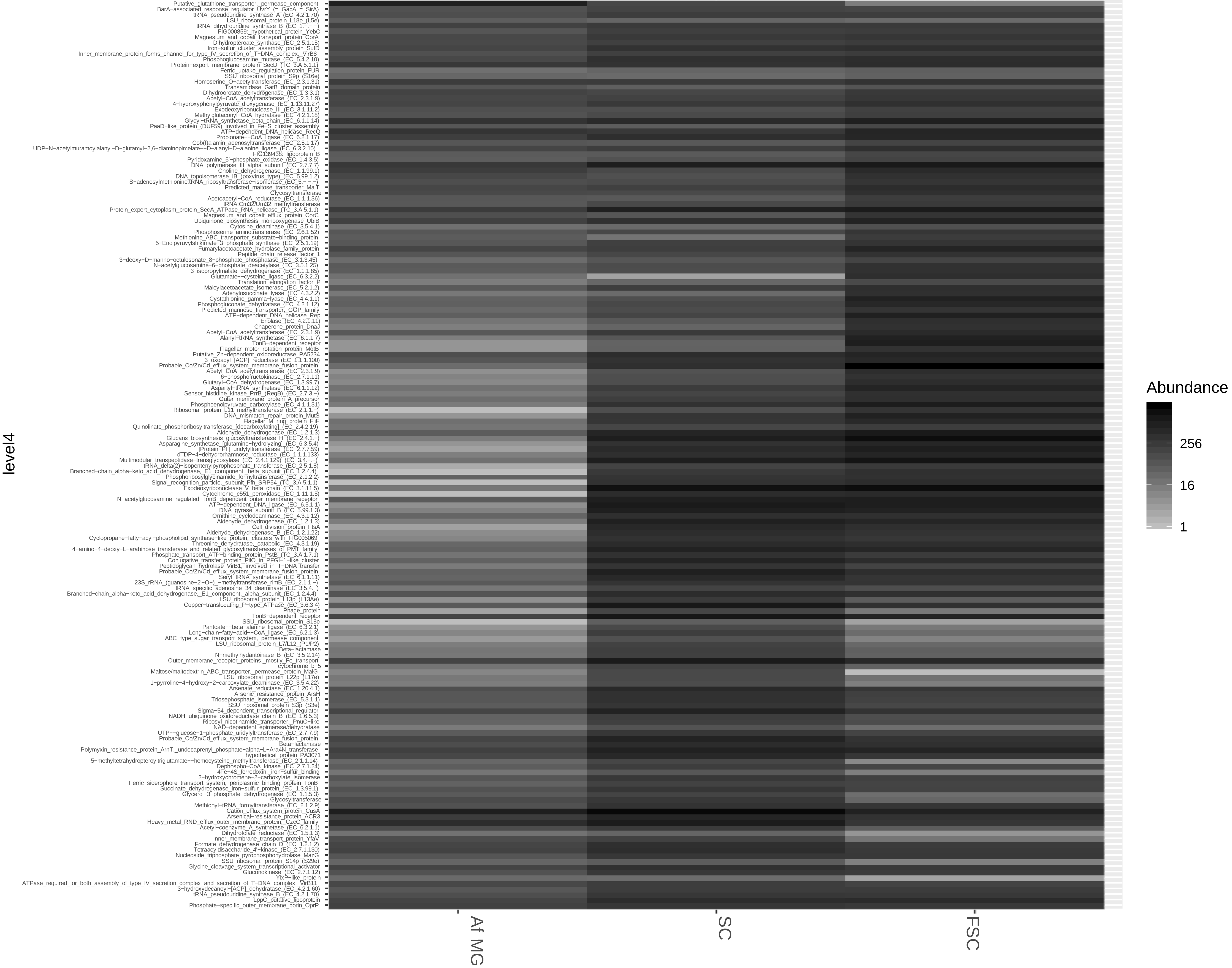


**Supplementary Figure S5. Abundances of the 166 shared annotated proteins.** Absolute frequency of the 166 proteins shared in the Af-MG, SC, and FSC Metagenomes, annotated with the level4 of the SEED Subsystems database.

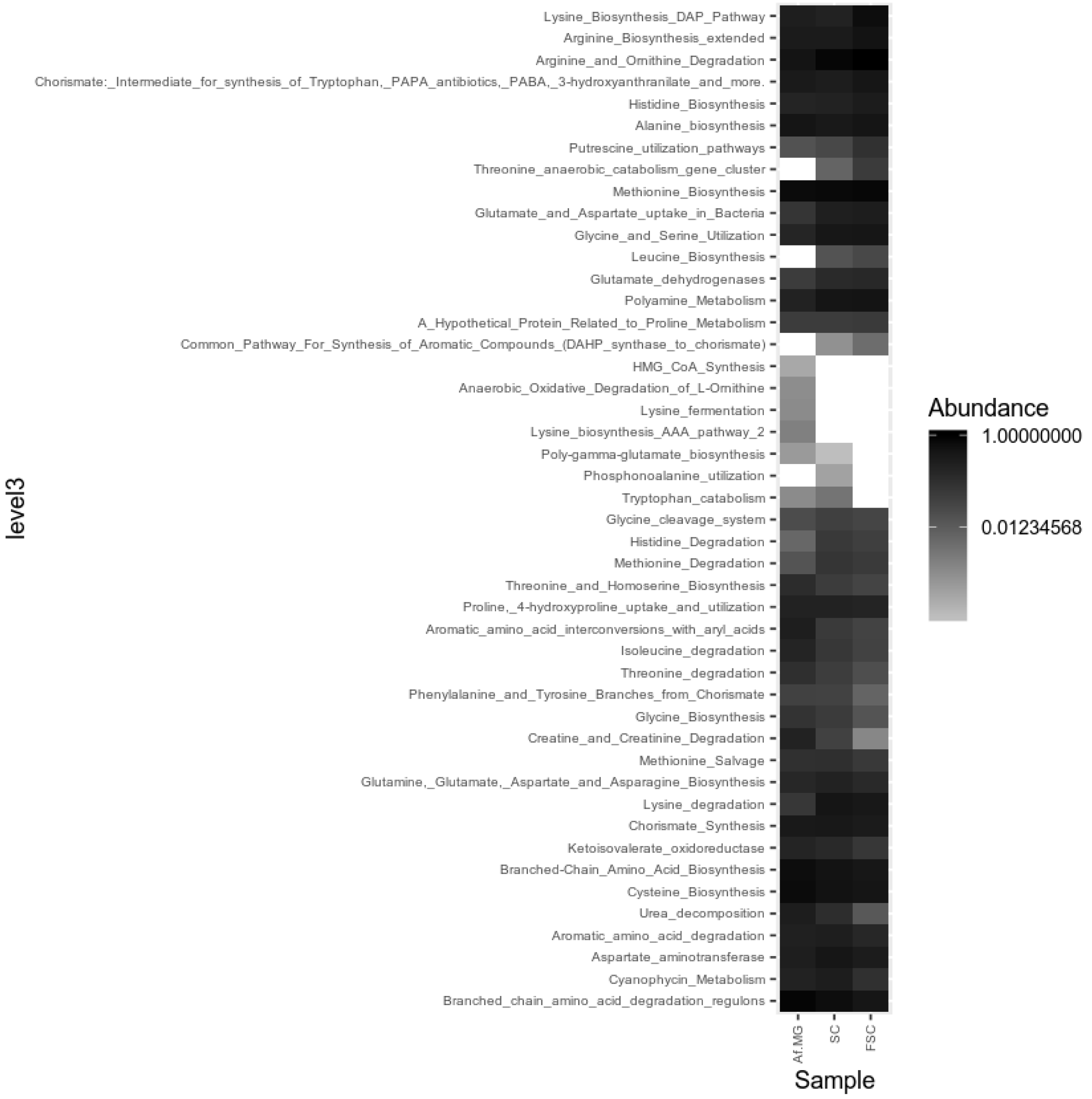


A

Amino Acids and Derivatives, level 3.

**Supplementary Figure S6. Relative abundance of SEED subsystems in the Af-MG, SC, and FSC metagenomes.** Heatmaps showingMapped read counts of each protein were normalized with the total read count for all proteins annotated with the subsystems database. Specific subsystems are shown: (A) Amino Acids and Derivatives, level 3; (B) Membrane transport, level 3; (C) Iron acquisition and metabolism, level 3; (D) Miscellaneous, level 3; (E) Clustering based subsystems, level 2; (F) Nitrogen metabolism, level 4; (G) Aromatic amino acid degradation (Amino acids and derivatives), level 4.

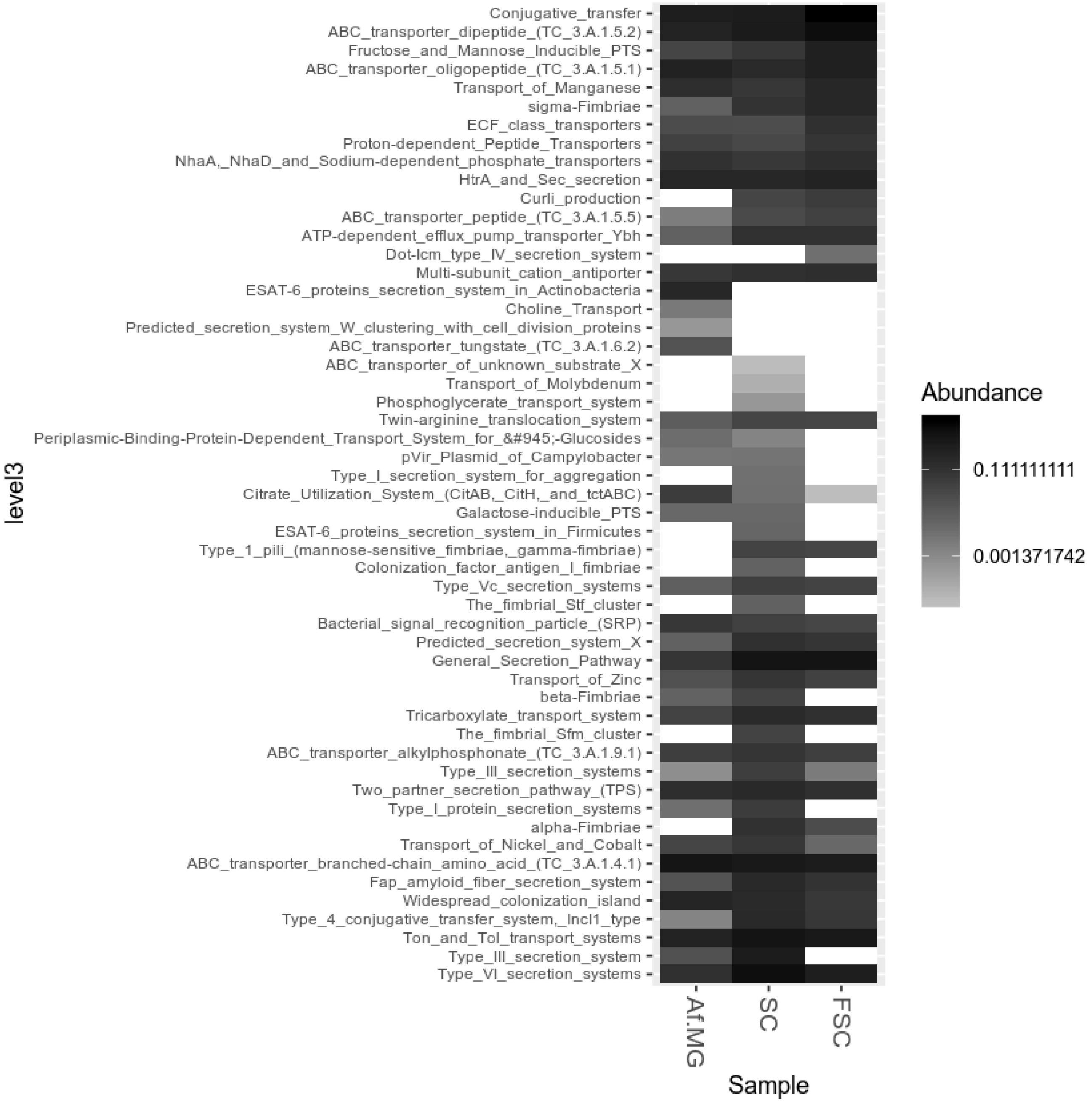


(B) Membrane transport, level 3.

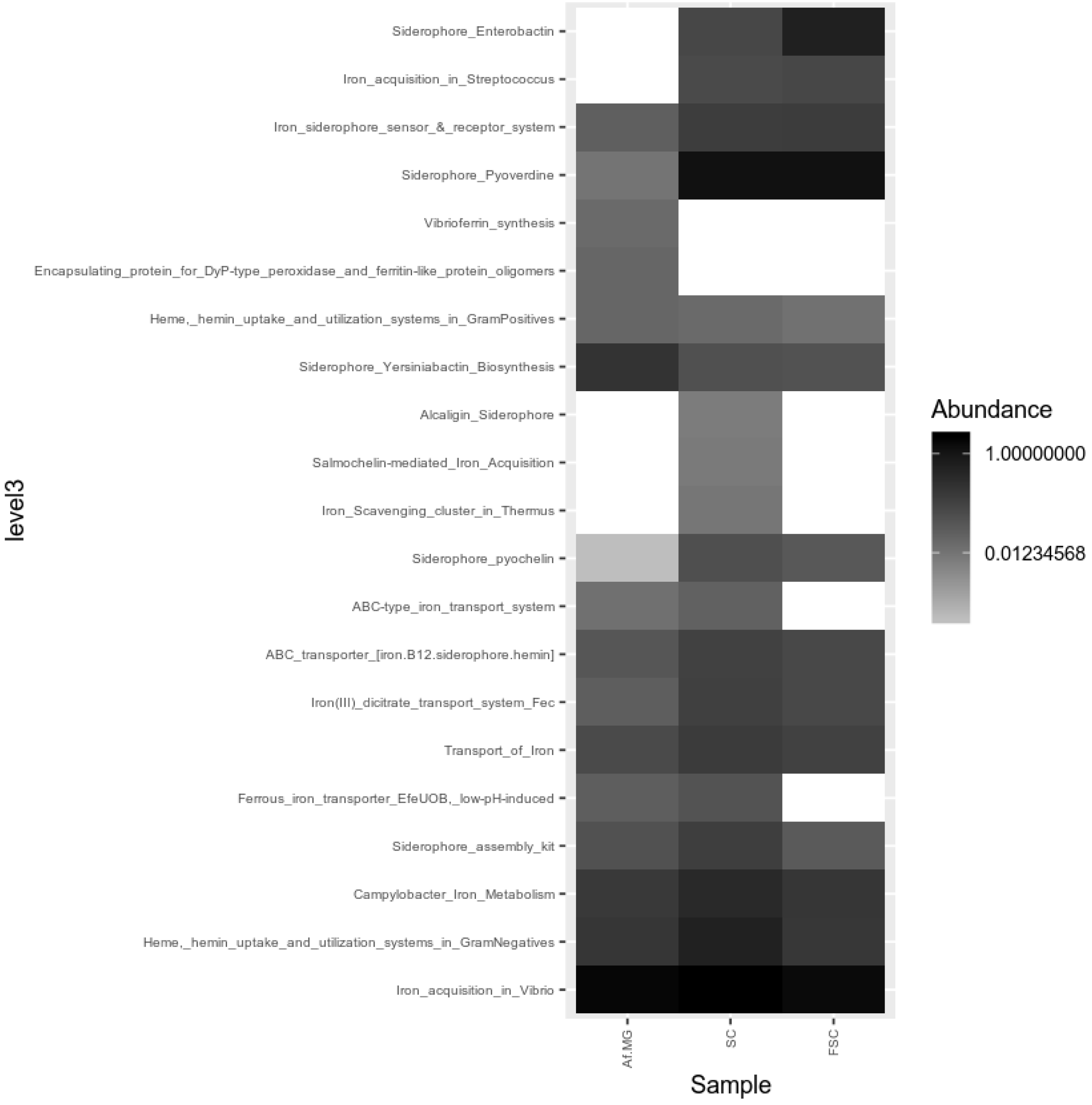


(C) Iron acquisition and metabolism, level 3.

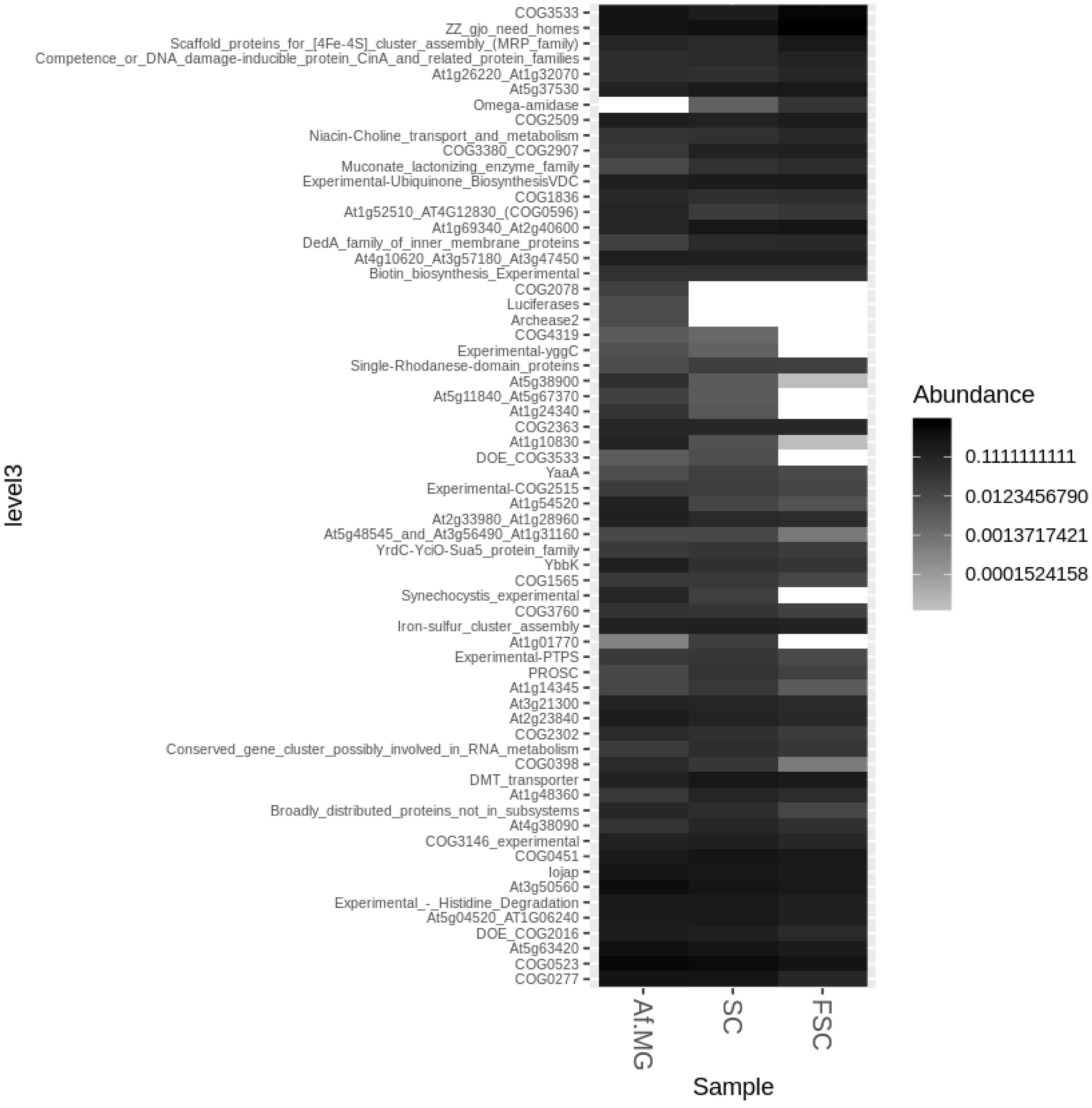


(D) Miscellaneous, level 3.

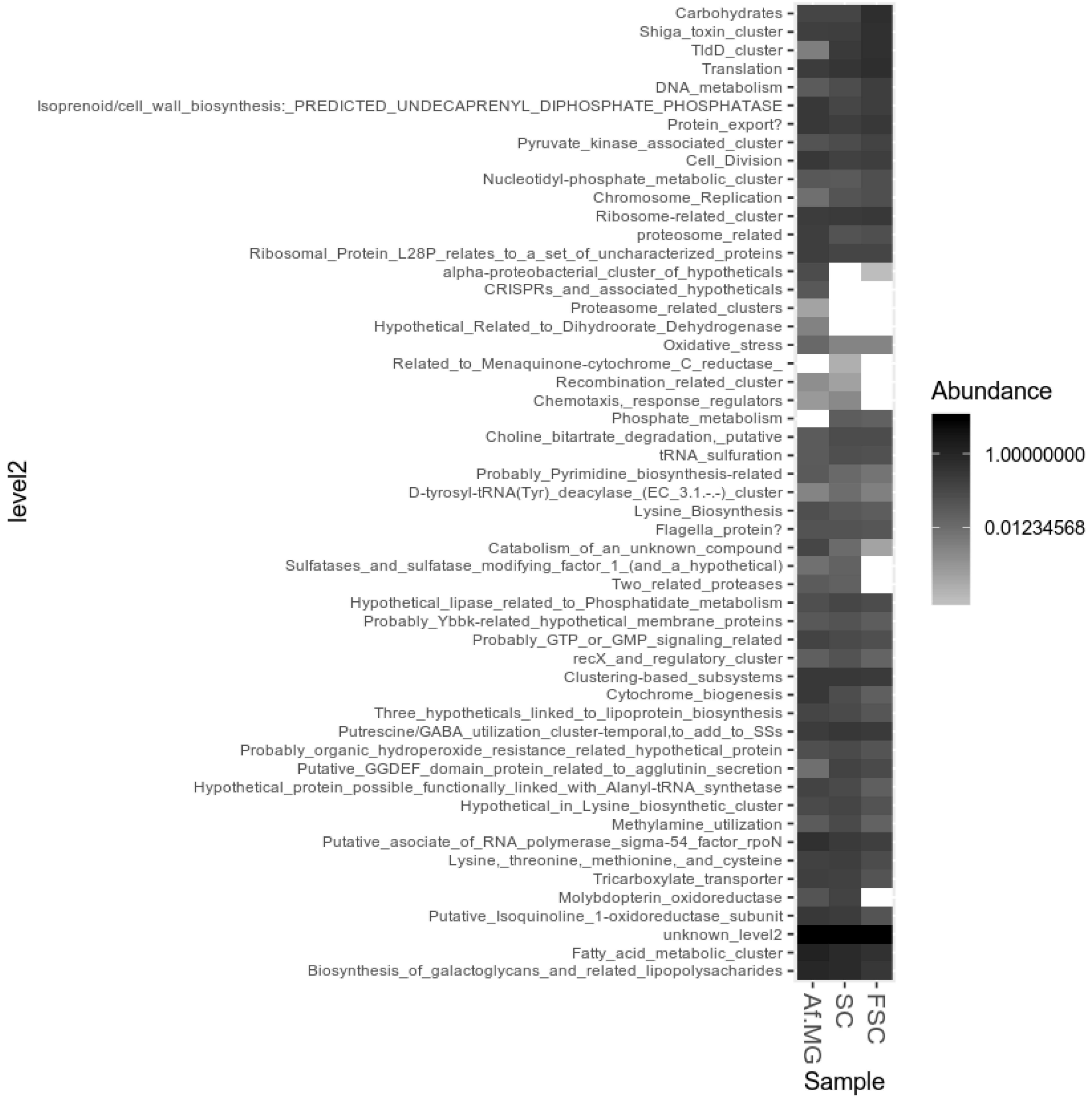


(E) Clustering based subsystems, level 2.

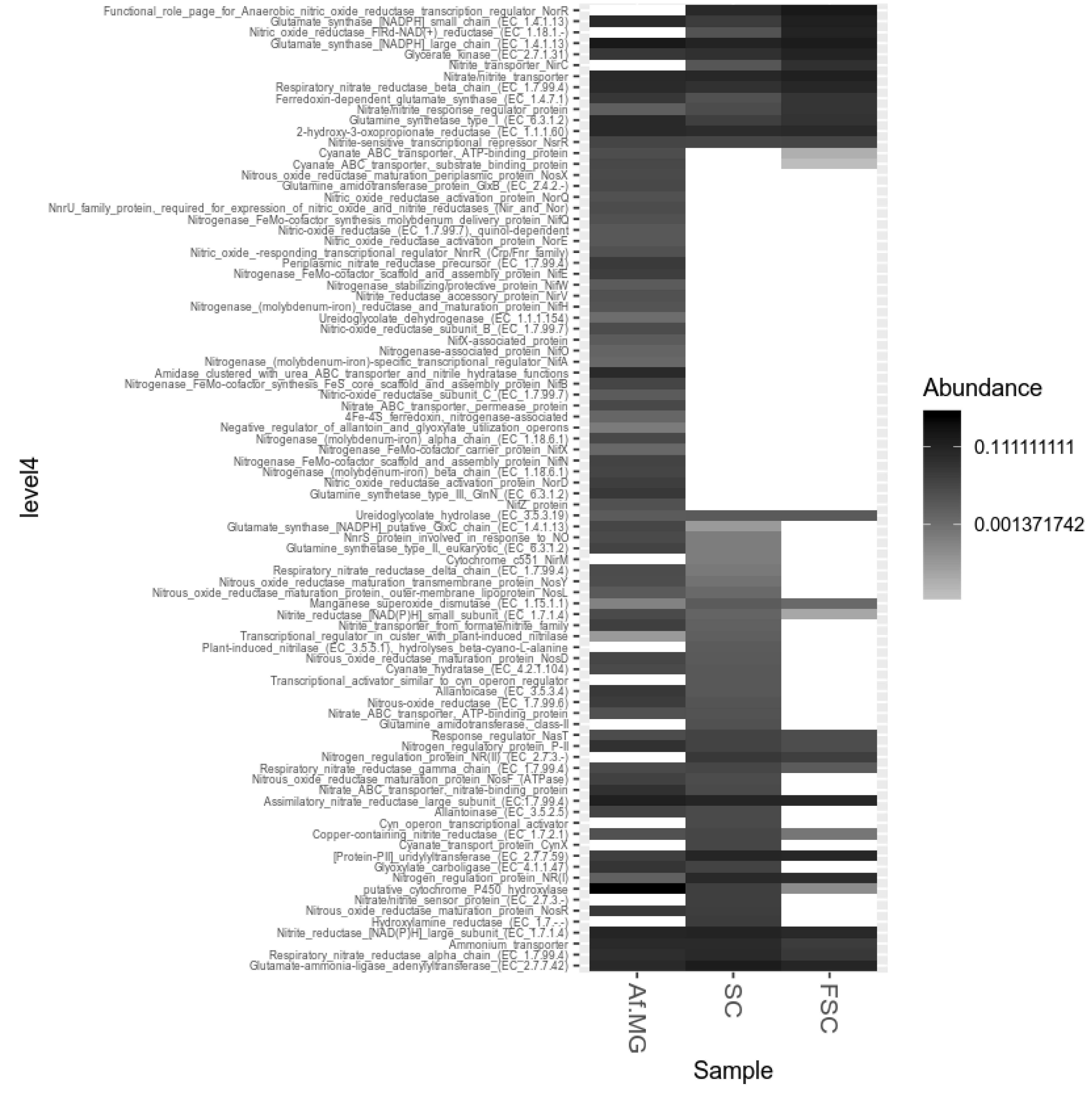


(F) Nitrogen metabolism, level 4.

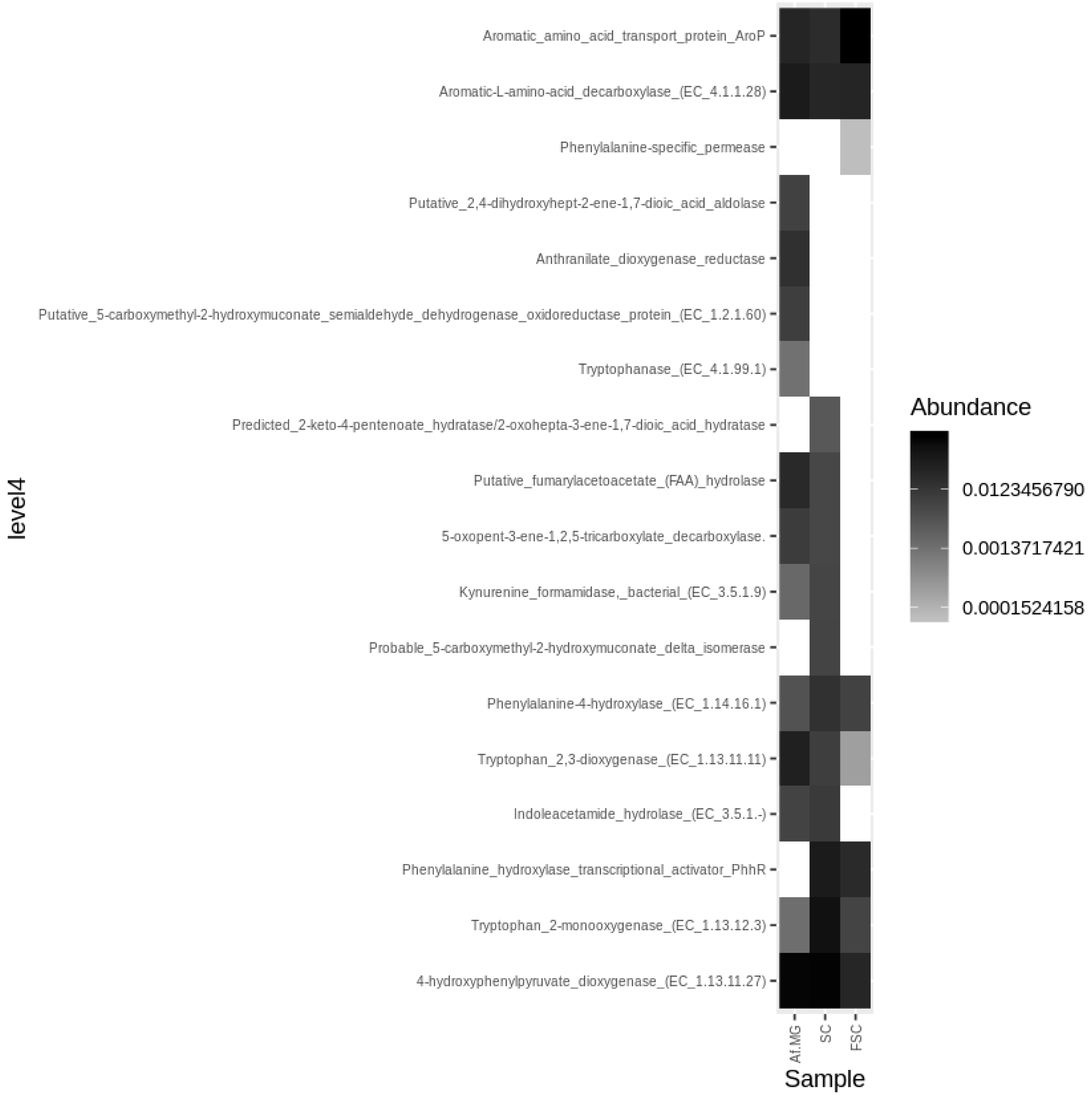


(G) Aromatic amino acid degradation (Amino acids and derivatives), level 4.

A

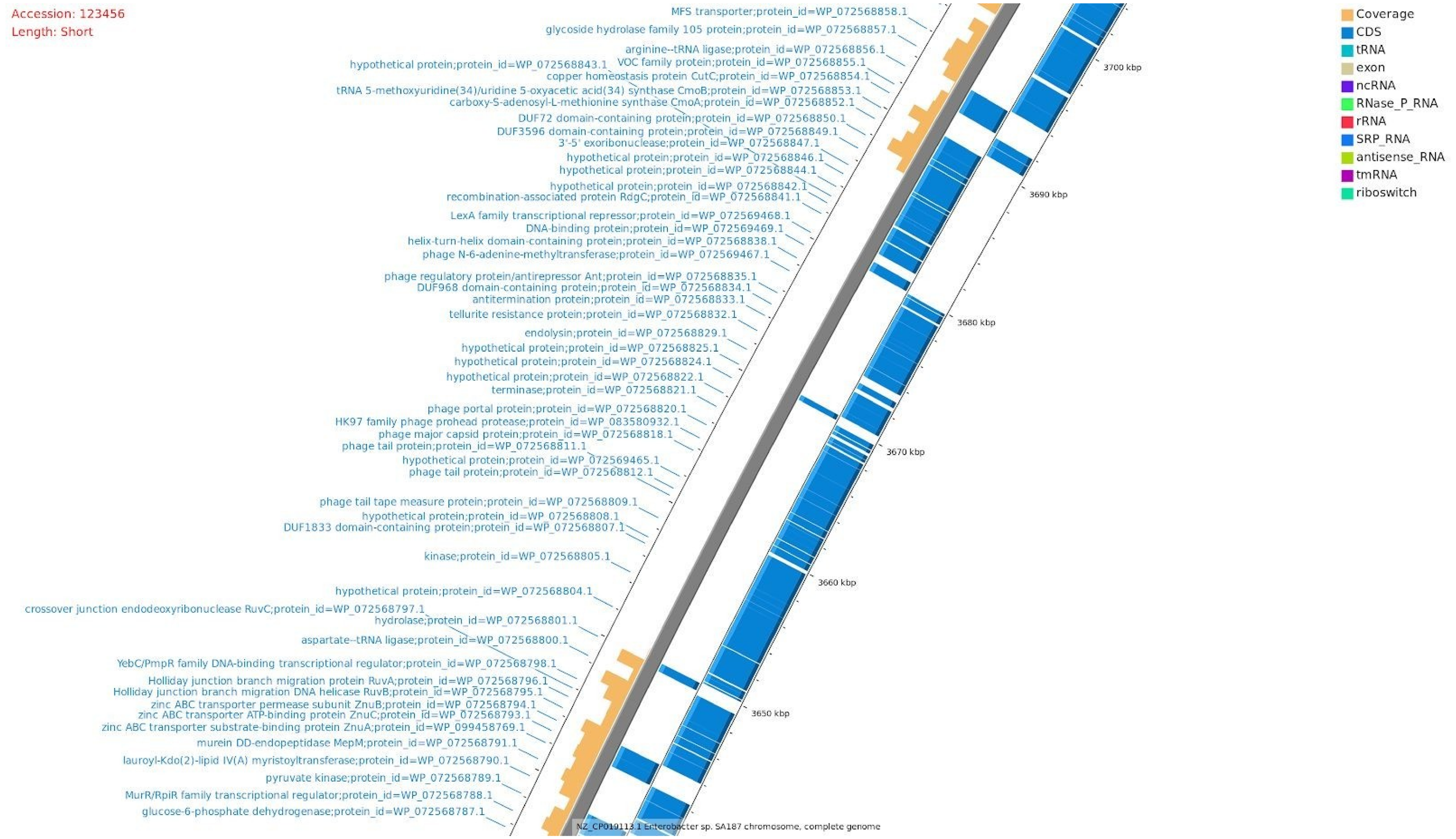


**Supplementary Figure S7. Unmapped loci of the *Enterobacter* sp. SA187 genome with the FSC metagenomic reads.** (A) 3,640 kbp – 3,700 kbp. (B) 2,200 kbp – 2,210 kbp. (C) 1,595 – 1,620 kbp. (D) 870 kbp – 900 kbp. (E) 440 kbp – 480 kbp. (F) 1,710 kbp – 1,760 kbp.

B

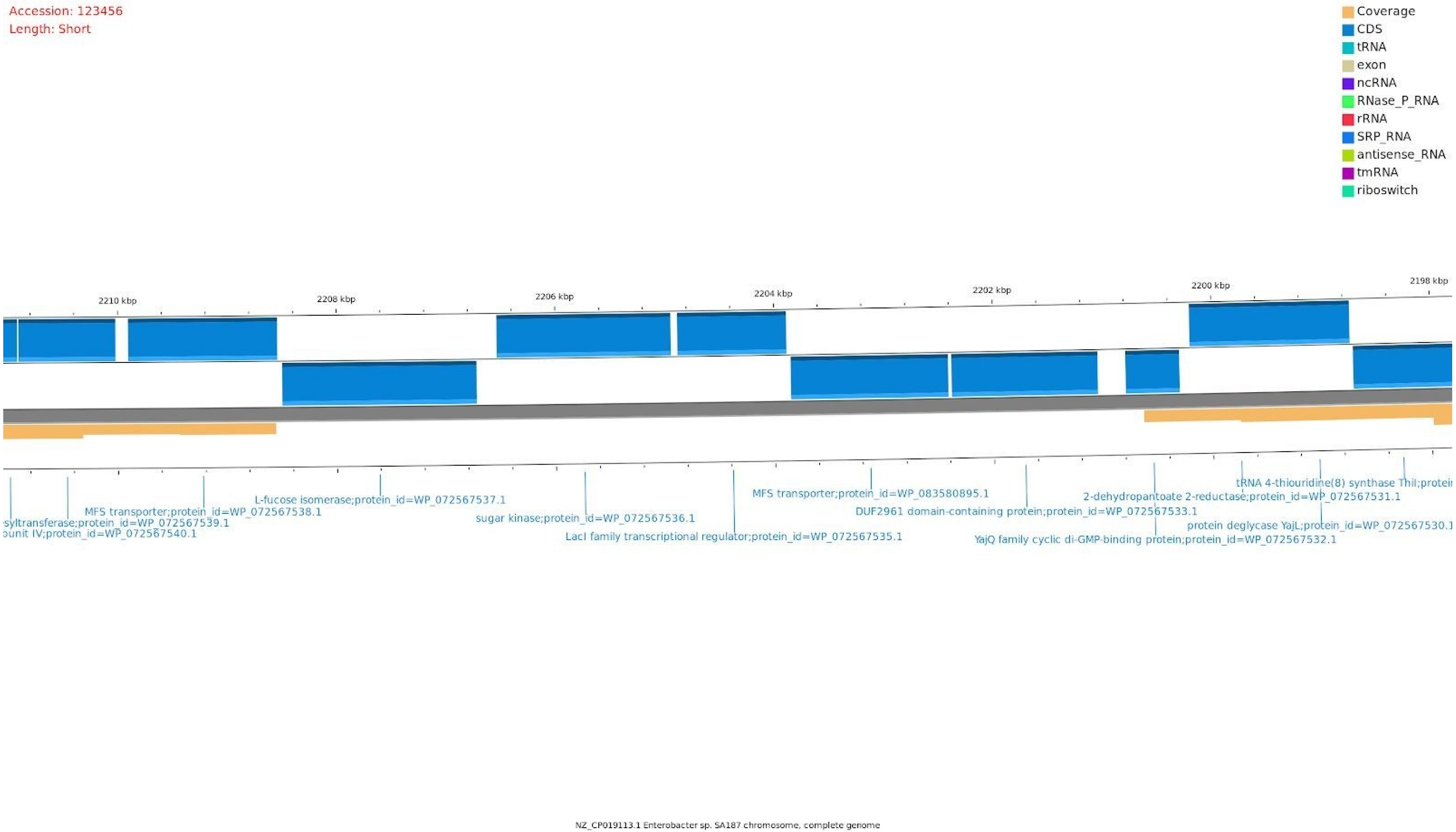
Accession: 123456
Length: Short
Coverage
CDS
tRNA
exon
ncRNA
RNase_P_RNA
rRNA
SRP_RNA
antisense_RNA
tmRNA
riboswitch
2210 kbp
2208 kbp
2206 kbp
2204 kbp
2202 kbp
2200 kbp
2198 kbp
L-fucose isomerase;protein_id=WP_072567537.1
MFS transporter;protein_id=WP_072567538.1
syltransferase;protein_id=WP_072567539.1
ounit IV;protein_id=WP_072567540.1
sugar kinase;protein_id=WP_072567536.1
LacI family transcriptional regulator;protein_id=WP_072567535.1
MFS transporter;protein_id=WP_083580895.1
DUF2961 domain-containing protein;protein_id=WP_072567533.1
YajQ family cyclic di-GMP-binding protein;protein_id=WP_072567532.1
2-dehydropantoate 2-reductase;protein_id=WP_072567531.1
tRNA 4-thiouridine(8) synthase ThiI;protei
protein deglycase YajL;protein_id=WP_072567530.1
NZ_CP019113.1 Enterobacter sp. SA187 chromosome, complete genome

C

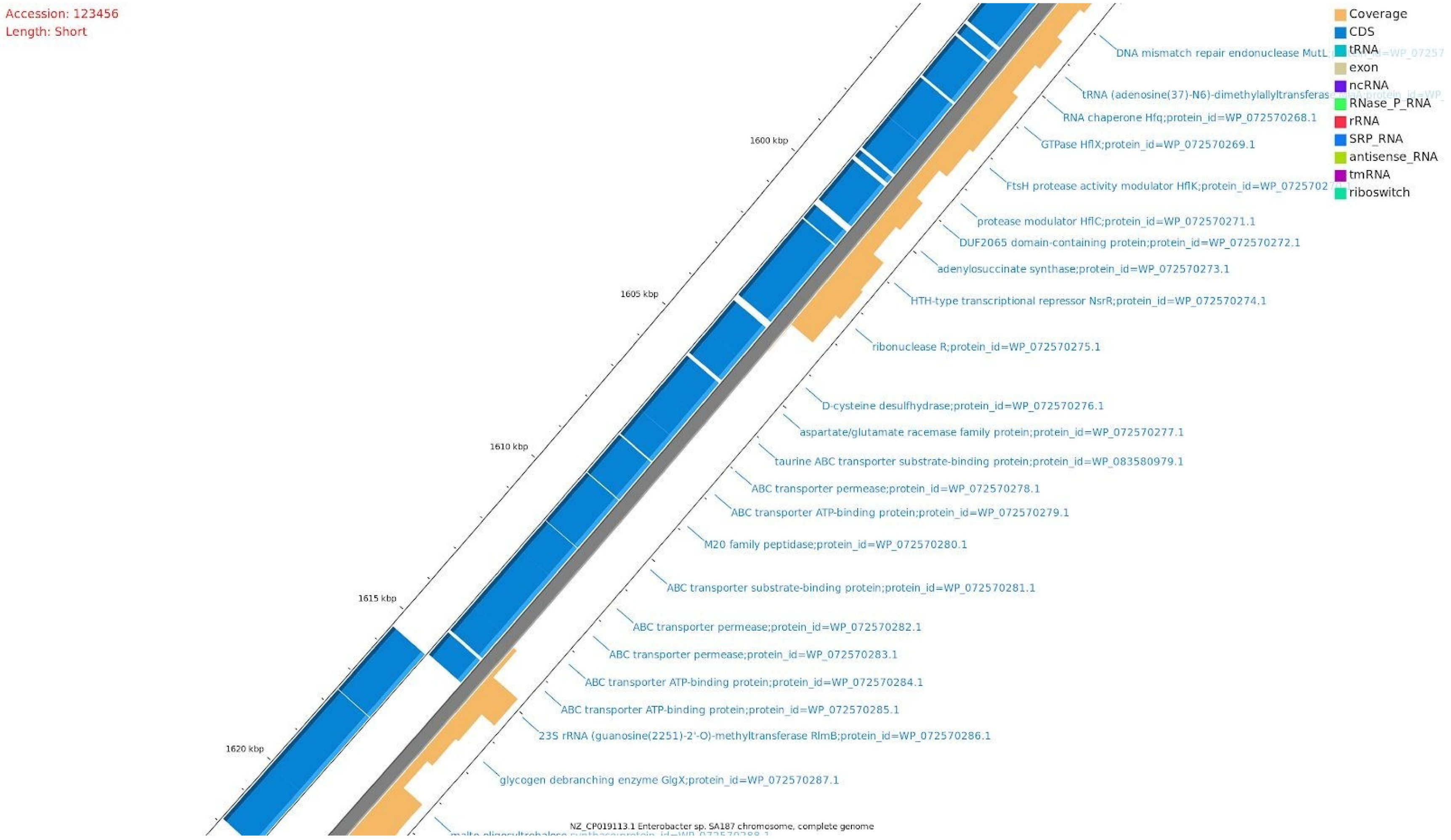

Accession: 123456
Length: Short
Coverage
CDS
tRNA
exon
ncRNA
RNase_P_RNA
rRNA
SRP_RNA
antisense_RNA
tmRNA
riboswitch
1600 kbp
1605 kbp
1610 kbp
1615 kbp
1620 kbp
DNA mismatch repair endonuclease MutL
tRNA (adenosine(37)-N6)-dimethylallyltransferas
RNA chaperone Hfq;protein_id=WP_072570268.1
GTPase HflX;protein_id=WP_072570269.1
FtsH protease activity modulator HflK;protein_id=WP_0725702
protease modulator HflC;protein_id=WP_072570271.1
DUF2065 domain-containing protein;protein_id=WP_072570272.1
adenylosuccinate synthase;protein_id=WP_072570273.1
HTH-type transcriptional repressor NsrR;protein_id=WP_072570274.1
ribonuclease R;protein_id=WP_072570275.1
D-cysteine desulfhydrase;protein_id=WP_072570276.1
aspartate/glutamate racemase family protein;protein_id=WP_072570277.1
taurine ABC transporter substrate-binding protein;protein_id=WP_083580979.1
ABC transporter permease;protein_id=WP_072570278.1
ABC transporter ATP-binding protein;protein_id=WP_072570279.1
M20 family peptidase;protein_id=WP_072570280.1
ABC transporter substrate-binding protein;protein_id=WP_072570281.1
ABC transporter permease;protein_id=WP_072570282.1
ABC transporter permease;protein_id=WP_072570283.1
ABC transporter ATP-binding protein;protein_id=WP_072570284.1
ABC transporter ATP-binding protein;protein_id=WP_072570285.1
23S rRNA (guanosine(2251)-2'-O)-methyltransferase RlmB;protein_id=WP_072570286.1
glycogen debranching enzyme GlgX;protein_id=WP_072570287.1
NZ_CP019113.1 Enterobacter sp. SA187 chromosome, complete genome

D

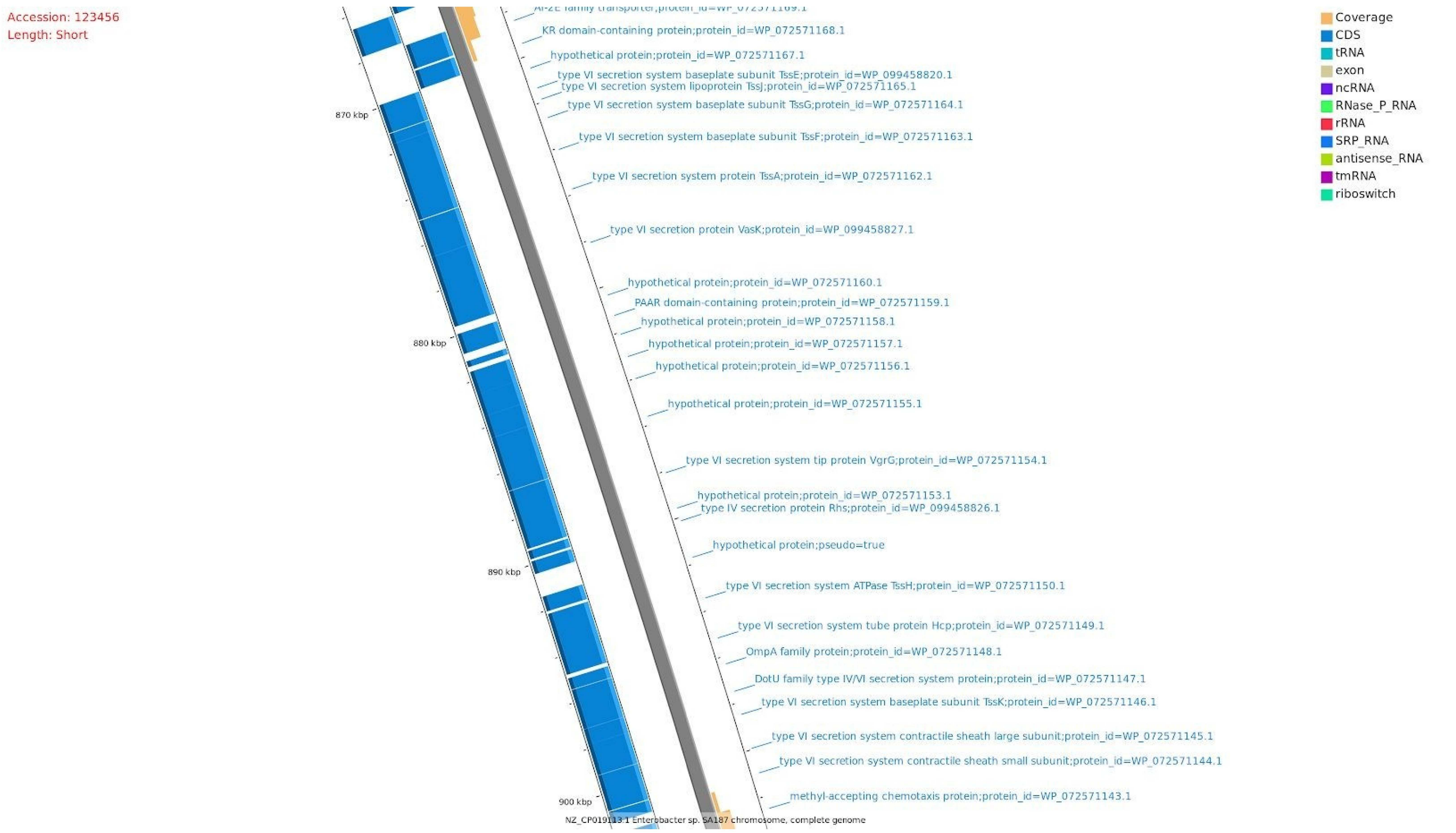

Accession: 123456
Length: Short
KR domain-containing protein;protein_id=WP_072571168.1
hypothetical protein;protein_id=WP_072571167.1
type VI secretion system baseplate subunit TssE;protein_id=WP_099458820.1
type VI secretion system lipoprotein TssJ;protein_id=WP_072571165.1
type VI secretion system baseplate subunit TssG;protein_id=WP_072571164.1
type VI secretion system baseplate subunit TssF;protein_id=WP_072571163.1
type VI secretion system protein TssA;protein_id=WP_072571162.1
type VI secretion protein VasK;protein_id=WP_099458827.1
hypothetical protein;protein_id=WP_072571160.1
PAAR domain-containing protein;protein_id=WP_072571159.1
hypothetical protein;protein_id=WP_072571158.1
hypothetical protein;protein_id=WP_072571157.1
hypothetical protein;protein_id=WP_072571156.1
hypothetical protein;protein_id=WP_072571155.1
type VI secretion system tip protein VgrG;protein_id=WP_072571154.1
hypothetical protein;protein_id=WP_072571153.1
type IV secretion protein Rhs;protein_id=WP_099458826.1
hypothetical protein;pseudo=true
type VI secretion system ATPase TssH;protein_id=WP_072571150.1
type VI secretion system tube protein Hcp;protein_id=WP_072571149.1
OmpA family protein;protein_id=WP_072571148.1
DotU family type IV/VI secretion system protein;protein_id=WP_072571147.1
type VI secretion system baseplate subunit TssK;protein_id=WP_072571146.1
type VI secretion system contractile sheath large subunit;protein_id=WP_072571145.1
type VI secretion system contractile sheath small subunit;protein_id=WP_072571144.1
methyl-accepting chemotaxis protein;protein_id=WP_072571143.1
870 kbp
880 kbp
890 kbp
900 kbp
NZ_CP019113.1 Enterobacter sp. SA187 chromosome, complete genome
Coverage
CDS
tRNA
exon
ncRNA
RNase_P_RNA
rRNA
SRP_RNA
antisense_RNA
tmRNA
riboswitch

F

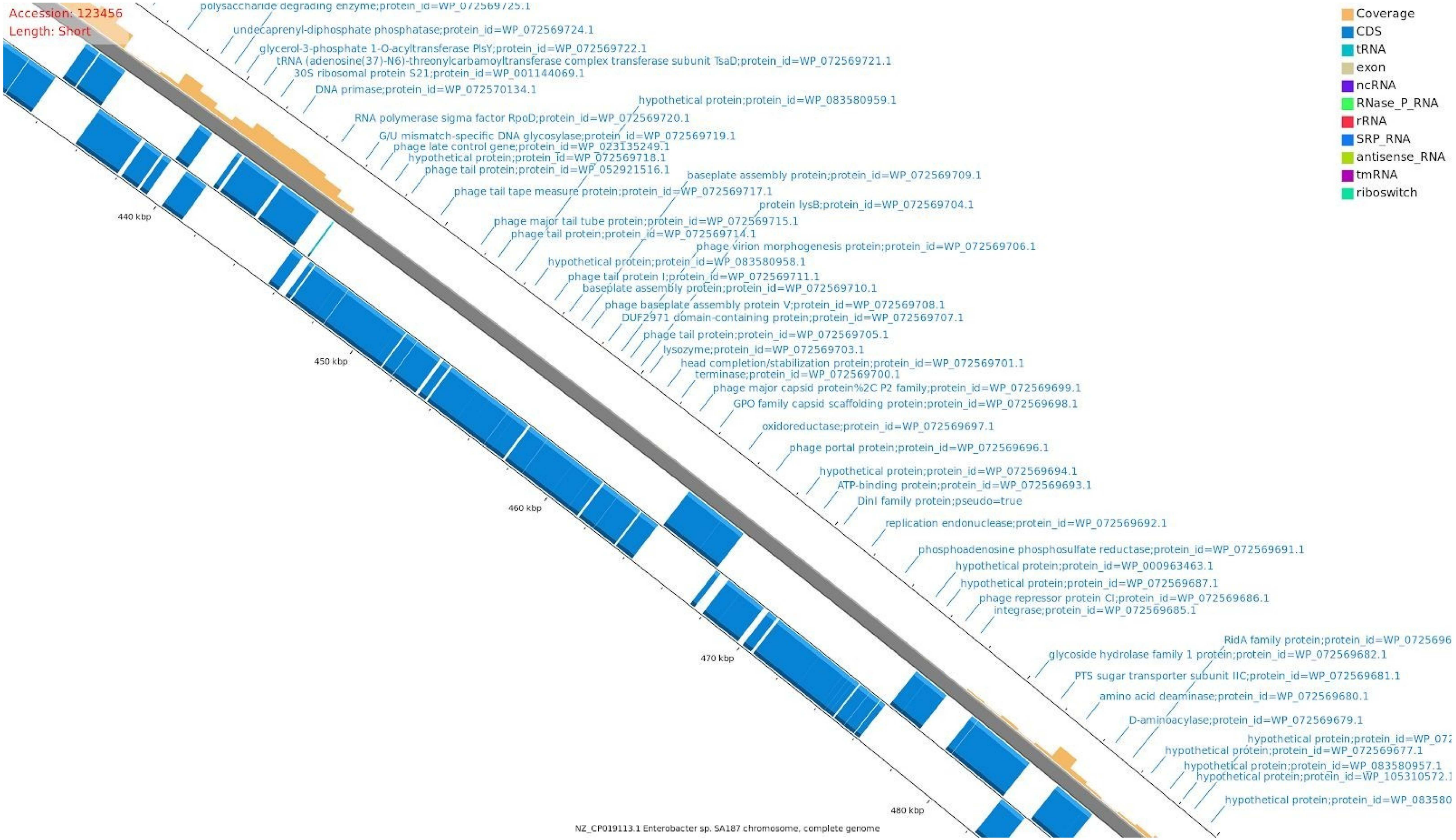

Accession: 123456
Length: Short
Coverage
CDS
tRNA
exon
ncRNA
RNase_P_RNA
rRNA
SRP_RNA
antisense_RNA
tmRNA
riboswitch
polysaccharide degrading enzyme;protein_id=WP_072569725.1
undecaprenyl-diphosphate phosphatase;protein_id=WP_072569724.1
glycerol-3-phosphate 1-O-acyltransferase PlsY;protein_id=WP_072569722.1
tRNA (adenosine(37)-N6)-threonylcarbamoyltransferase complex transferase subunit TsaD;protein_id=WP_072569721.1
30S ribosomal protein S21;protein_id=WP_001144069.1
DNA primase;protein_id=WP_072570134.1
hypothetical protein;protein_id=WP_083580959.1
RNA polymerase sigma factor RpoD;protein_id=WP_072569720.1
G/U mismatch-specific DNA glycosylase;protein_id=WP_072569719.1
phage late control gene;protein_id=WP_023135249.1
hypothetical protein;protein_id=WP_072569718.1
phage tail protein;protein_id=WP_052921516.1
baseplate assembly protein;protein_id=WP_072569709.1
phage tail tape measure protein;protein_id=WP_072569717.1
protein lysB;protein_id=WP_072569704.1
phage major tail tube protein;protein_id=WP_072569715.1
phage tail protein;protein_id=WP_072569714.1
phage virion morphogenesis protein;protein_id=WP_072569706.1
hypothetical protein;protein_id=WP_083580958.1
phage tail protein I;protein_id=WP_072569711.1
baseplate assembly protein;protein_id=WP_072569710.1
phage baseplate assembly protein V;protein_id=WP_072569708.1
DUF2971 domain-containing protein;protein_id=WP_072569707.1
phage tail protein;protein_id=WP_072569705.1
lysozyme;protein_id=WP_072569703.1
head completion/stabilization protein;protein_id=WP_072569701.1
terminase;protein_id=WP_072569700.1
phage major capsid protein%2C P2 family;protein_id=WP_072569699.1
GPO family capsid scaffolding protein;protein_id=WP_072569698.1
oxidoreductase;protein_id=WP_072569697.1
phage portal protein;protein_id=WP_072569696.1
hypothetical protein;protein_id=WP_072569694.1
ATP-binding protein;protein_id=WP_072569693.1
DinI family protein;pseudo=true
replication endonuclease;protein_id=WP_072569692.1
phosphoadenosine phosphosulfate reductase;protein_id=WP_072569691.1
hypothetical protein;protein_id=WP_000963463.1
hypothetical protein;protein_id=WP_072569687.1
phage repressor protein CI;protein_id=WP_072569686.1
integrase;protein_id=WP_072569685.1
RidA family protein;protein_id=WP_0725696
glycoside hydrolase family 1 protein;protein_id=WP_072569682.1
PTS sugar transporter subunit IIC;protein_id=WP_072569681.1
amino acid deaminase;protein_id=WP_072569680.1
D-aminoacylase;protein_id=WP_072569679.1
hypothetical protein;protein_id=WP_072
hypothetical protein;protein_id=WP_072569677.1
hypothetical protein;protein_id=WP_083580957.1
hypothetical protein;protein_id=WP_105310572.1
hypothetical protein;protein_id=WP_083580
440 kbp
450 kbp
460 kbp
470 kbp
480 kbp
NZ_CP019113.1 Enterobacter sp. SA187 chromosome, complete genome

G

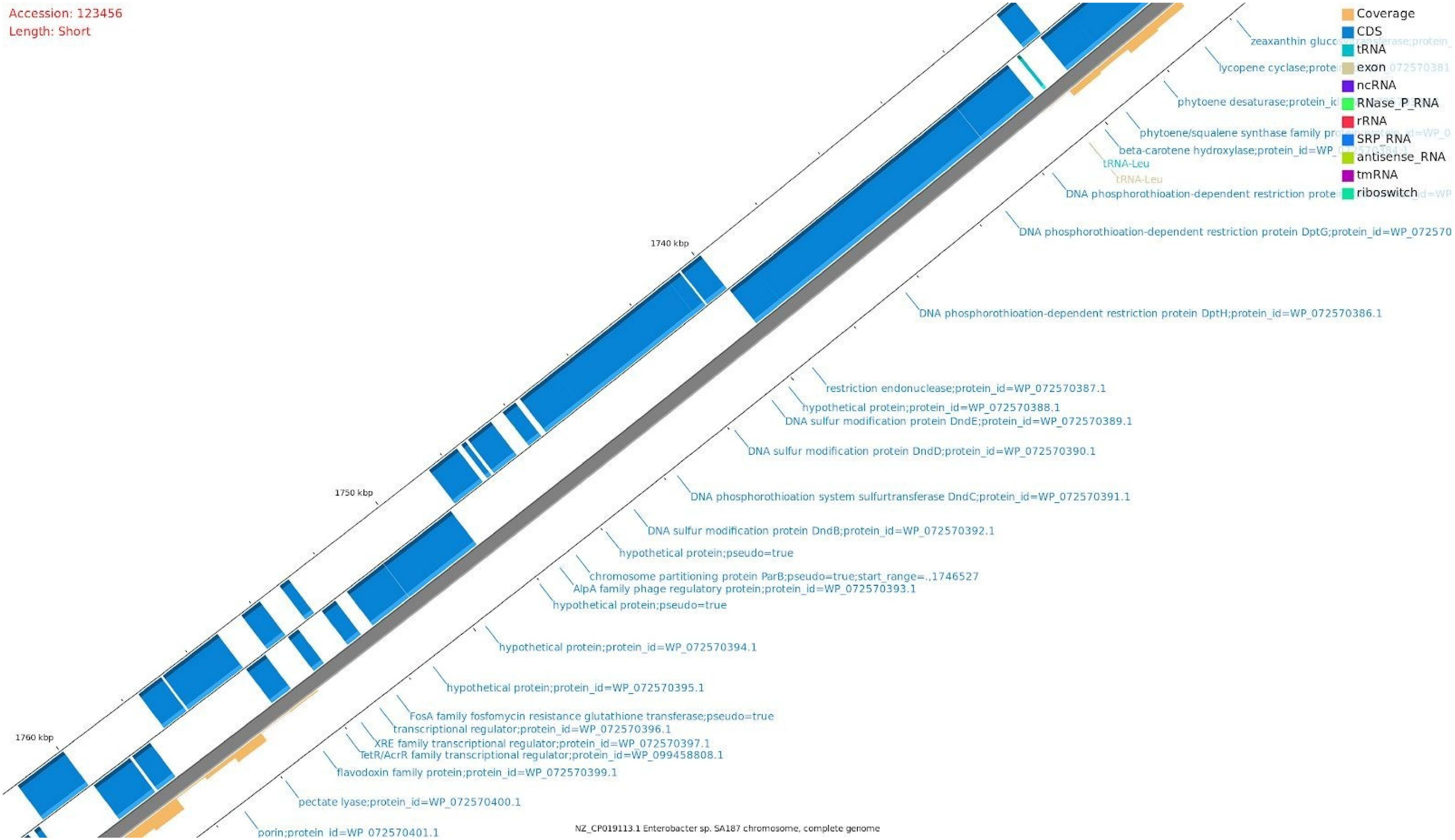
Accession: 123456
Length: Short
Coverage
CDS
tRNA
exon
ncRNA
RNase_P_RNA
rRNA
SRP_RNA
antisense_RNA
tmRNA
riboswitch
1740 kbp
1750 kbp
1760 kbp
lycopene cyclase;prote
phytoene desaturase;protein_id
phytoene/squalene synthase family pr
beta-carotene hydroxylase;protein_id=WP_
tRNA-Leu
tRNA-Leu
DNA phosphorothioation-dependent restriction prote
DNA phosphorothioation-dependent restriction protein DptG;protein_id=WP_072570
DNA phosphorothioation-dependent restriction protein DptH;protein_id=WP_072570386.1
restriction endonuclease;protein_id=WP_072570387.1
hypothetical protein;protein_id=WP_072570388.1
DNA sulfur modification protein DndE;protein_id=WP_072570389.1
DNA sulfur modification protein DndD;protein_id=WP_072570390.1
DNA phosphorothioation system sulfurtransferase DndC;protein_id=WP_072570391.1
DNA sulfur modification protein DndB;protein_id=WP_072570392.1
hypothetical protein;pseudo=true
chromosome partitioning protein ParB;pseudo=true;start_range=.,1746527
AlpA family phage regulatory protein;protein_id=WP_072570393.1
hypothetical protein;pseudo=true
hypothetical protein;protein_id=WP_072570394.1
hypothetical protein;protein_id=WP_072570395.1
FosA family fosfomycin resistance glutathione transferase;pseudo=true
transcriptional regulator;protein_id=WP_072570396.1
XRE family transcriptional regulator;protein_id=WP_072570397.1
TetR/AcrR family transcriptional regulator;protein_id=WP_099458808.1
flavodoxin family protein;protein_id=WP_072570399.1
pectate lyase;protein_id=WP_072570400.1
porin;protein_id=WP_072570401.1
NZ_CP019113.1 Enterobacter sp. SA187 chromosome, complete genome

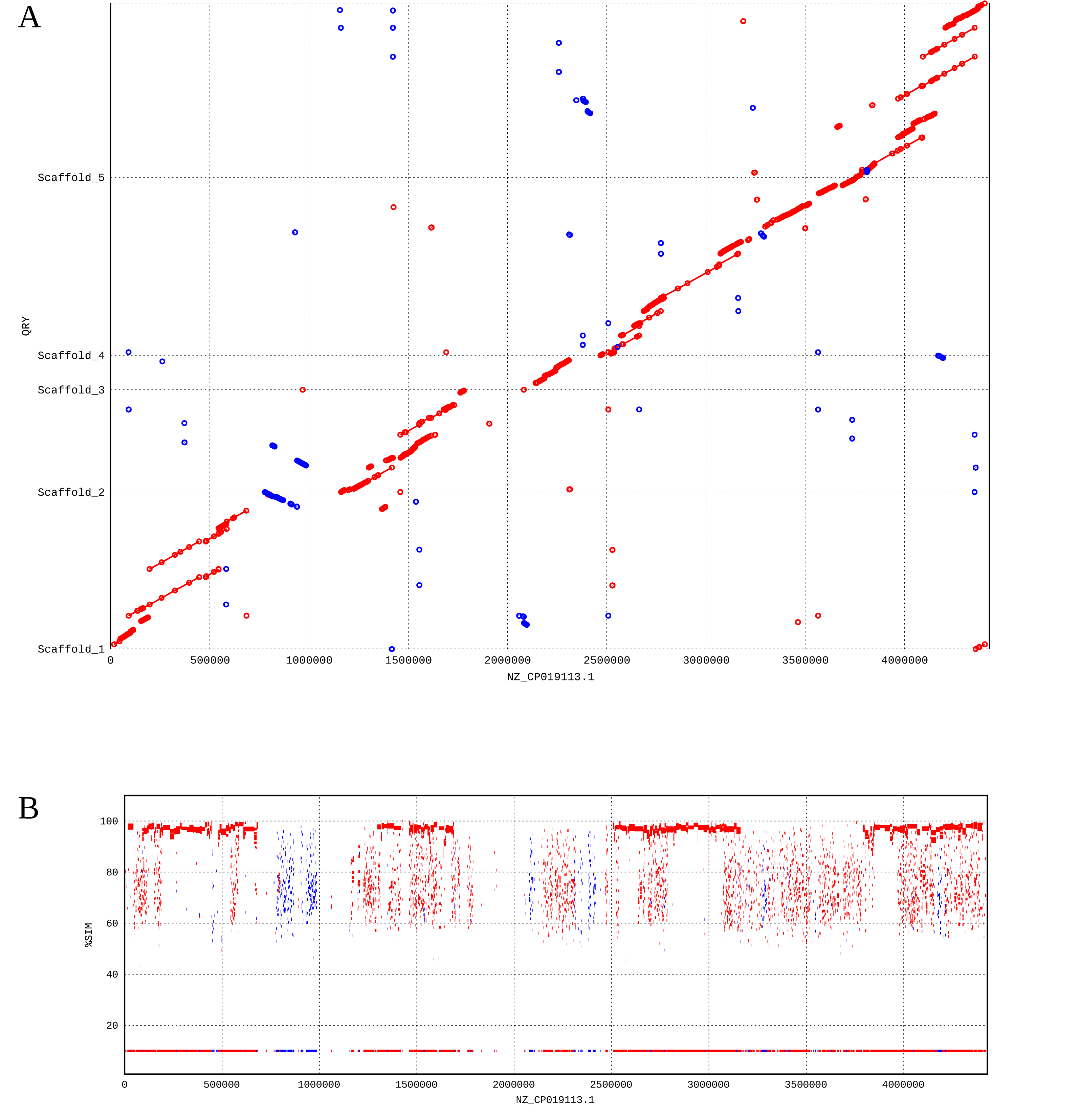


**Supplementary Figure S8. Comparison of the assembled scaffolds from the metagenomic data to the genome of *Enterobacter* sp. SA187.** Alignment of nucleotides (A) and predicted protein sequences (B) of the scaffolds built with the metagenomic contigs assigned to the *Enterobacter* genus with Kraken and ordered with scaffold_builder.

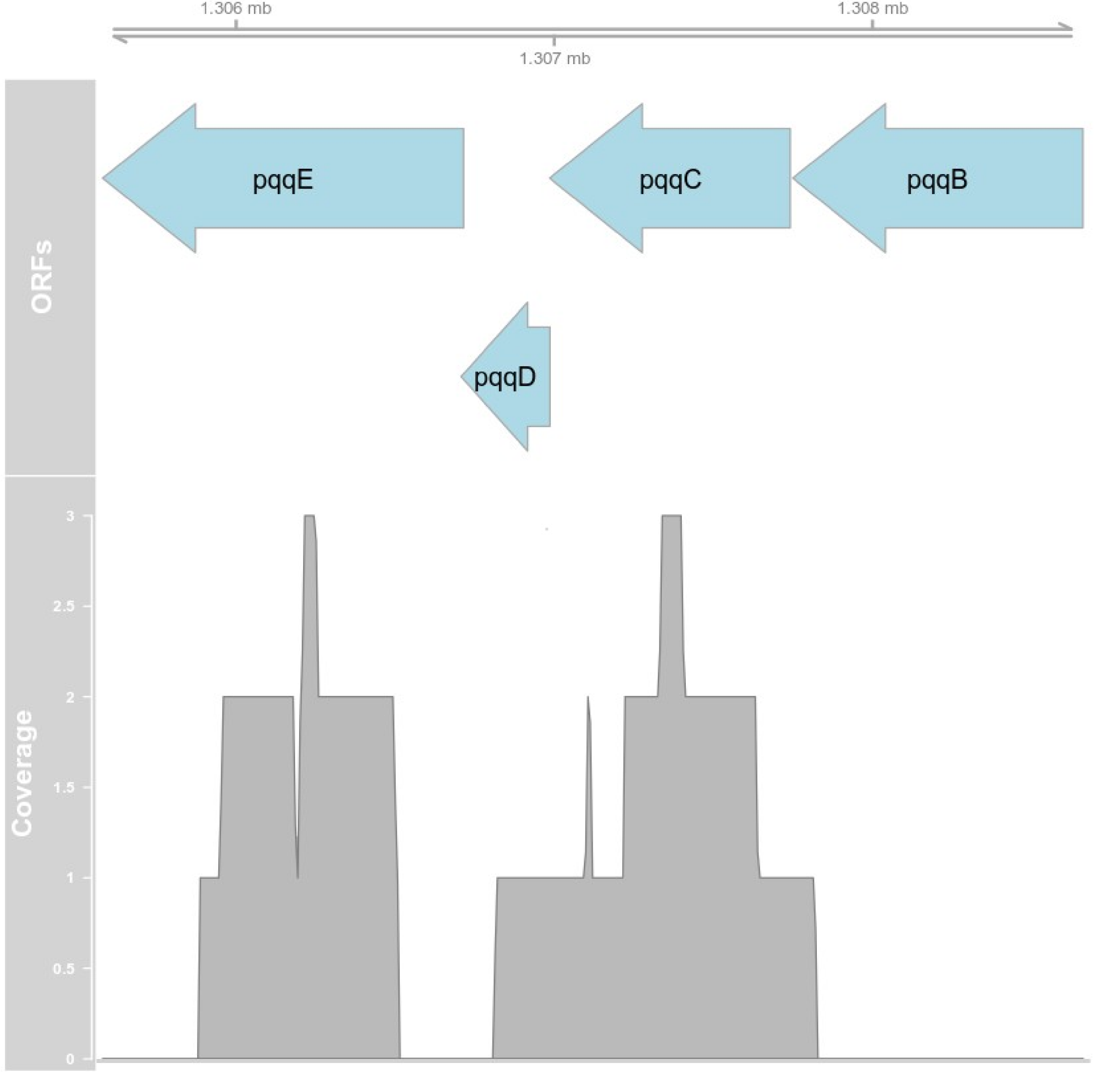


**Supplementary Figure S9.** Metagenomic read mapping of the pqqBCDE genes in the built partial genome. Locus of the pqqBCDE genes in the Scaffold 1 of the partially assembled genome. ORFs are shown as arrows. The bottom track shows the aligned reads with Bowtie2 against the scaffold.
